# RIP1 kinase mediates angiogenesis by modulating macrophages in experimental neovascularization

**DOI:** 10.1101/739599

**Authors:** Takashi Ueta, Kenji Ishihara, Shoji Notomi, Jong-Jer Lee, Daniel Maidana, Nikolaos Efstathiou, Yusuke Murakami, Eiichi Hasegawa, Kunihiro Azuma, Tetsuya Toyono, Eleftherios Paschalis, Makoto Aihara, Joan W. Miller, Demetrios G. Vavvas

## Abstract

Inflammation plays an important role in pathologic angiogenesis. Receptor-interacting protein 1 (RIP1) is highly expressed in inflammatory cells and is known to play an important role in the regulation of apoptosis, necroptosis, and inflammation, however its role in angiogenesis remains elusive. Here, we show that RIP1 is abundantly expressed in infiltrating macrophages during angiogenesis, and genetic or pharmacological inhibition of RIP1 kinase activity using kinase-inactive RIP1^K45A/K45A^ mice or necrostatin-1 attenuates angiogenesis in laser-induced choroidal neovascularization (CNV), Matrigel plug angiogenesis, and alkali injury-induced corneal neovascularization in mice. The inhibitory effect on angiogenesis was mediated by caspase activation through a kinase-independent function of RIP1 and RIP3, and simultaneous caspase inhibition with RIP1 kinase inhibition abrogated the effects of RIP1 kinase inhibition on angiogenesis *in vivo*. Mechanistically, infiltrating macrophages are the key target for RIP1 kinase inhibition to attenuate pathological angiogenesis, and we observed that the inhibition of RIP1 kinase activity is associated with caspase activation in infiltrating macrophages and decreased expression of pro-angiogenic M2-like markers while M1 marker expressions were sustained. Similarly, *in vitro*, catalytic inhibition of RIP1 down-regulated M2 marker expressions in IL-4-activated bone marrow-derived macrophages, which was blocked by simultaneous caspase inhibition. Taken together, these results suggest a novel, non-necrotic function of RIP1 kinase activity and suggest that RIP1-mediated modulation of macrophage activation may represent a therapeutic target for the control of angiogenesis-related diseases.

**Significance:** Pathological angiogenesis has been implicated in diverse pathologies. Infiltrating macrophages, especially those activated to M2-like phenotype are critically important to support angiogenesis. Whereas the role of RIP1 kinase in the regulation of apoptosis, necroptosis, and inflammation have been well established, its role in angiogenesis remains elusive despite being abundantly expressed in angiogenesis-related infiltrating macrophages. This study demonstrated for the first time that RIP1 kinase inhibition attenuates angiogenesis in multiple mouse models of pathological angiogenesis *in vivo*. Mechanistically, the inhibitory effect on angiogenesis depends on RIP kinase inhibition-mediated caspase activation in infiltrating macrophages that suppresses M2-like polarization, thereby attenuating pathological angiogenesis.

## INTRODUCTION

Pathologic angiogenesis has been implicated in many disorders, including neovascular age-related macular degeneration (AMD), the leading cause of blindness in the elderly worldwide.(1) During pathological angiogenesis, the abundant infiltration of macrophages supports vascular endothelial sprouting. Tissue-infiltrating macrophages are polarized into functional phenotypes, including classically activated (pro-inflammatory M1) and alternatively activated (anti-inflammatory and tissue-remodeling M2) macrophages. Macrophages can undergo dynamic transitions between different functional states, potentially driven by the tissue microenvironment that dictates a transcriptional response.(2) In angiogenesis, the importance of M2-like macrophages or the M1-to-M2 transition of phenotypes has been well established through several angiogenesis models,(3–8) including laser-induced choroidal neovascularization (CNV), a major mouse model of neovascular AMD.(3,5,6,8) However, the current knowledge of the regulation of macrophage phenotypes and functions remains incomplete.

Previous studies have suggested a causative role for death receptors(9–13) and toll-like receptors (TLRs)(14–16) in angiogenesis, including neovascular AMD(10,13) and the well established model of laser-induced CNV in mice,(11,16) although a consensus has not been reached.(17,18) Receptor-interacting protein 1 (RIP1) is an intracellular adaptor protein that relays signals from these receptors to regulate inflammation, apoptosis and necroptosis.(19–22) RIP1 is subjected to posttranslational modifications, including ubiquitination, phosphorylation of the kinase domains, and oligomerization via interaction with RIP homotypic interaction motif (RHIM). Ubiquitinated RIP1 induces NF-κB-mediated inflammation and cell survival, whereas deubiquitinated RIP1 forms a ‘ripoptosome’ with caspase-8, cellular FLICE-inhibitory protein (cFLIP), and Fas-associated death domain (FADD), leading to either apoptosis or necroptosis.(19) Phosphorylated and RHIM-oligomerized RIP1 exerts catalytic kinase activity to form a ‘necrosome’ with RIP3 and mixed-lineage kinase-like protein, leading to necroptosis and inflammation.(19,20,22) Because caspase-8 cleaves RIP1 to inhibit the necroptotic pathway, apoptosis is frequently induced as the primary mode of cell death, while necroptosis can be facilitated when the activity of caspase-8 or other apoptotic factors, including FADD is down-regulated. Although kinase-dependent functions of RIP1 are usually pro-necrotic, kinase-independent functions of RIP1 could be important for preventing inappropriate activation of both caspase-8-dependent apoptosis and RIP3-dependent necroptosis.(23–26) The functions of RIP1 depend on cell types and the biological context, and its role in angiogenesis remains largely unknown despite its relative abundant expression in infiltrating macrophages.

In the present study, using genetically engineered mice and catalytic inhibitors for RIP1, RIP3, and caspases, we dissected a pivotal non-necrotic role of RIP1 in angiogenesis and macrophage polarization using *in vivo* and *in vitro* models of pathological angiogenesis.

## RESULTS

### Inhibition of RIP1 kinase activity suppresses CNV

To examine the role of RIP1 kinase activity in angiogenesis, we utilized laser-induced choroidal neovascularization (CNV) model, a well established and one of the most commonly used angiogenesis models (27). In this model there is upregulation of RIP kinases and caspases in retinal pigment epithelium (RPE)-choroid-sclera tissue containing CNV lesions during the phase of ongoing neovascular response with macrophage infiltration (Fig.1A) indicating a potential activation of the respective pathways. In addition, the same RPE-choroid tissue samples were dissected to the areas with CNV and those without CNV, and similar upregulation in RIP kinase protein levels were observed (See SI Appendix, Fig. S1). Immunohistochemistry also showed abundant expression of RIP1 protein in the laser-induced CNV lesion in mice (Fig. 1B). Furthermore, in human neovascular AMD, immunohistochemistry showed a relatively higher RIP1 protein expression in CD68-positive macrophages (Fig. 1C).

**Fig. 1.**
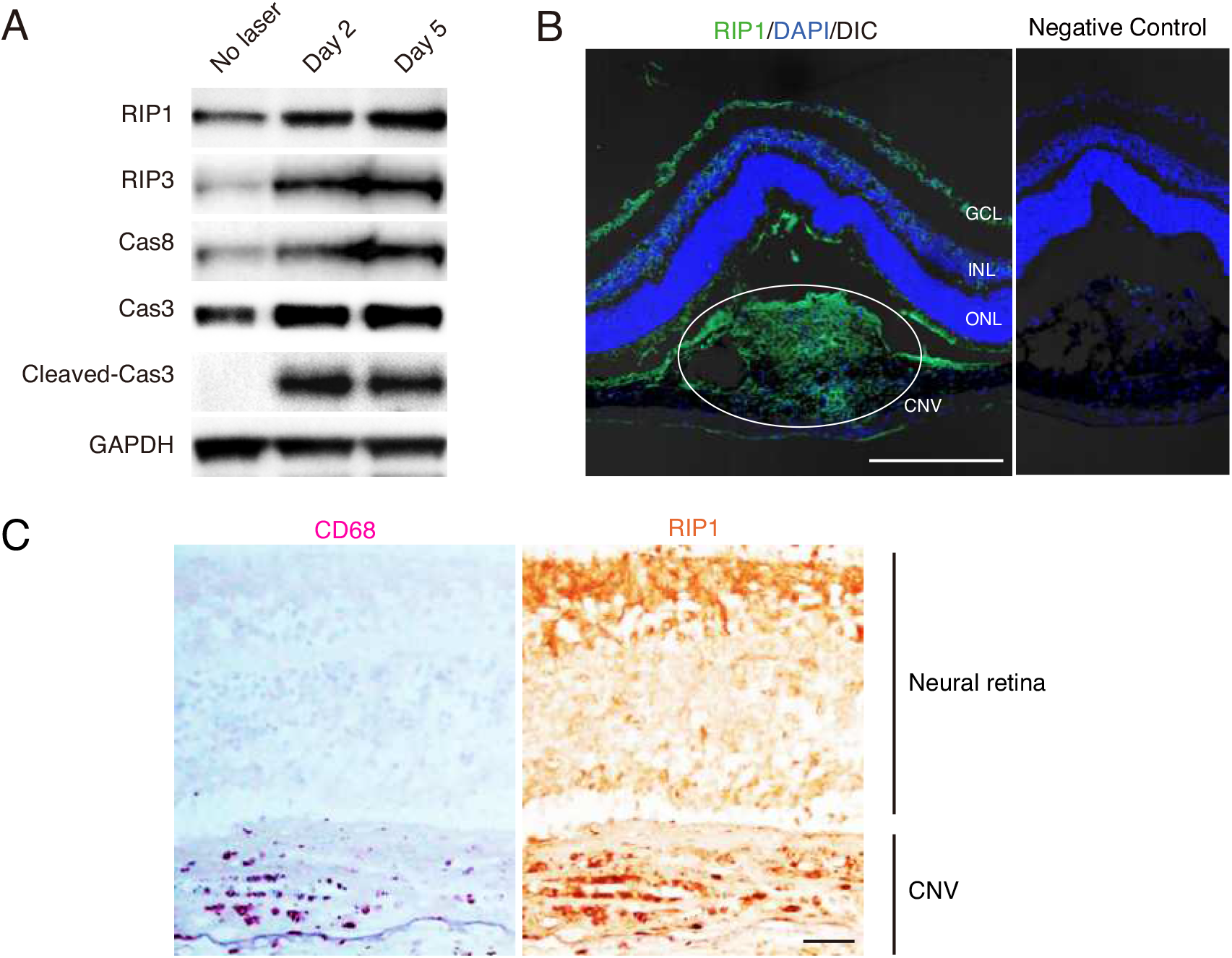
RIP1 expression in CNV. (A) Western blot analysis of RIP and caspase pathways in RPE-choroid on day 0, 2, and 5 after CNV induction. (B) Immunohistochemistry of RIP1 and its localization in F4/80(+) macrophages and CD31(+) endothelial cells in CNV on day 5 after induction by laser. Scale bar, 100 μm. GCL; ganglion cell layer, INL; inner nuclear layer, and ONL; outer nuclear layer of the retina. (C) Immunohistochemistry of RIP1 and CD68 in human neovascular AMD. Scale bar, 100μm.

We next examined the effect of the catalytic inhibition of RIP1 on the development of laser-induced CNV using necrostatin-1 (Nec-1). Nec-1 was administered through intravitreal injections immediately after CNV induction. We observed a dose-dependent effect of Nec-1 to reduce CNV lesion size (Fig. 2A). We also tested the effect of the systemic administration of Nec-1. Alzet osmotic pumps loaded with DMSO vehicle or Nec-1 were subcutaneously implanted the day before CNV induction. After 7 days of laser induction, Nec-1-treated mice had significantly smaller CNV sizes compared to control mice (Fig. 2B), confirming that Nec-1 effects are independent of administration procedures. Moreover, to rule out the possibility that Nec-1 effect on CNV is due to its influence on the acute cell death after laser irradiation, Nec-1 was intravitreally injected on day 4, which also significantly reduced the CNV size on day 7 (Fig. 2C). To address the longer-term effect of RIP1 kinase inhibition on CNV development, the effect of Nec-1 intravitreally injected on day 0 remained effective at least until day 14 (Fig. 2D). Lastly, we tested the additive effect of Nec-1 with the conventional anti-VEGF therapy using aflibercept. Combined aflibercept and Nec-1 treatment further reduced CNV size compared to aflibercept treatment alone (Fig. 2E).

**Fig. 2.**
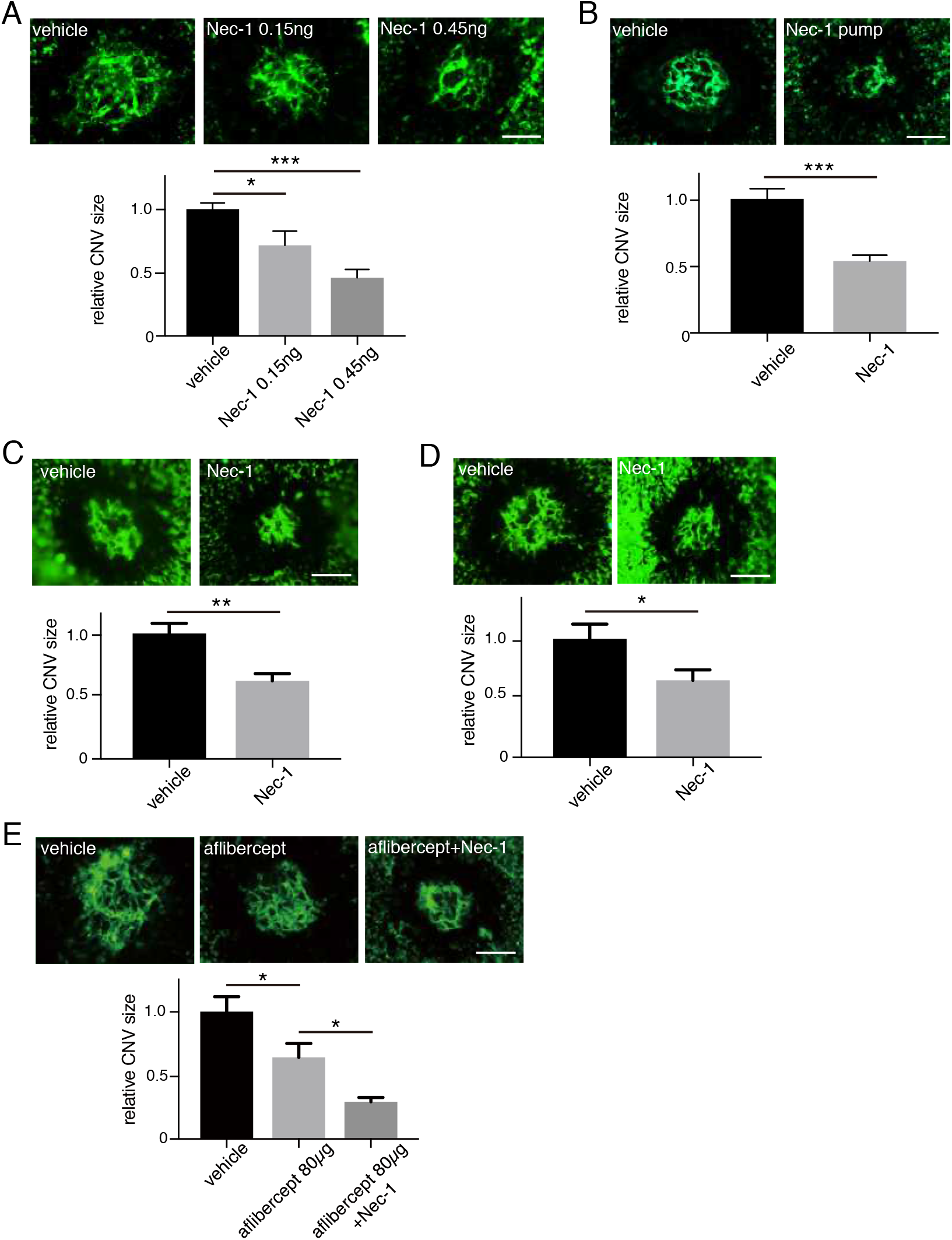
Nec-1 attenuates angiogenesis in laser-induced CNV. (A) After CNV induction (day 0), DMSO vehicle or Nec-1 was intravitreally injected to the eyes of WT mice, and CNV size was assessed on RPE-choroidal flat-mounts on day 7. n = 10–15 eyes per group. (B) DMSO vehicle or Nec-1 was administered through osmotic pumps that were subcutaneously implanted in WT mice the day before CNV induction. CNV size was assessed on RPE-choroidal flat-mounts on day 7. n = 6 eyes per group. (C) DMSO vehicle or Nec-1 was intravitreally injected to the eyes of WT mice on day 4 and CNV size was assesed on day 7. n=10 eyes per group. (D) DMSO vehicle or Nec-1 was intravitreally injected to the eyes of WT mice on day 3 and CNV size was assesed on day 14. n=10 eyes per group. (E) DMSO vehicle or Nec-1 was intravitreally injected to the eyes of WT mice on day 0, PBS vehicle or aflibercept was intravitreally injected on day 4, and CNV size was assesed on day 7. n=10 eyes per group. Scale bars, 100 μm. *P < 0.05, **P < 0.01, ***P < 0.001; Student’s t-test or one-way ANOVA and post-hoc Tukey’s test. Data are mean ± SEM.

### Caspase activation mediates the effect of RIP1 kinase inhibition to suppress CNV

To elucidate how the catalytic inhibition of RIP1 suppresses CNV, we first evaluated the involvement of RIP3 using RIP3-deficient mice and GSK’872, a catalytic inhibitor of RIP3. First, we observed that Nec-1 (RIP1 kinase inhibition) does not reduce CNV size in RIP3-deficient mice (Fig. 3A), indicating that RIP3 is necessary for Nec-1 to suppress CNV. Expectedly, the catalytic inhibition of RIP3 by GSK’872 decreased CNV size on day 7 in WT mice (Fig. 3B), while GSK’872 did not have an effect on RIP3-deficient mice (See SI Appendix, Fig. S2A). These results indicate that catalytic inhibition of RIP1 or RIP3 suppressed CNV, however, complete loss of RIP3 protein did not (See SI Appendix, Fig. S2B), suggesting a role for RIP scaffold in angiogenesis inhibition when the kinase activity is lost. We thus speculated that RIP kinase inhibition unmasks a kinase-independent RIP scaffold function that leads to CNV suppression.

**Fig. 3.**
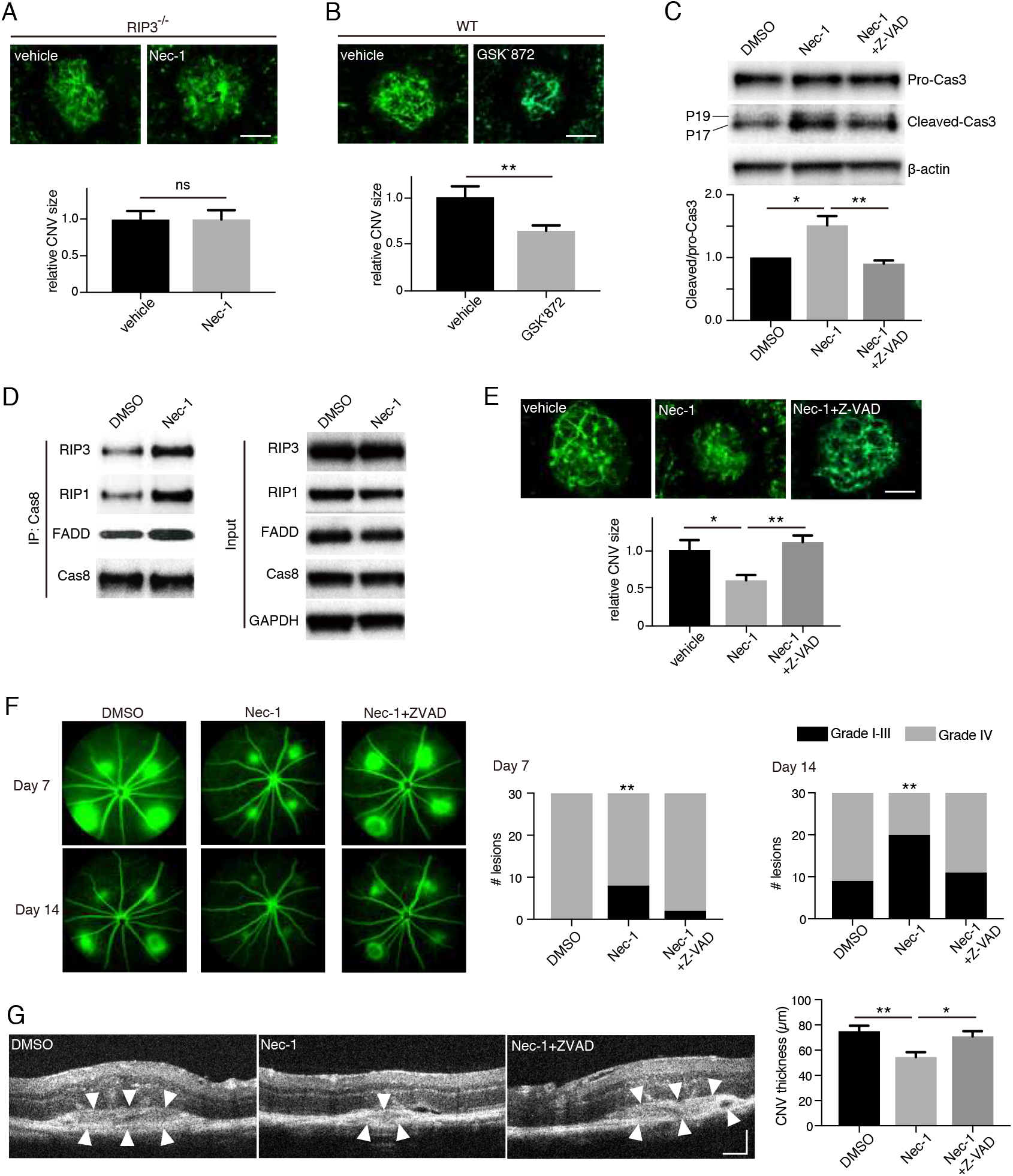
Caspase activation mediates the attenuation of CNV by catalytic inhibition of RIP1. (A) CNV size on flat-mounts on day 7 in RIP3−/− mice intravitreally injected with DMSO vehicle or Nec-1. n = 8 eyes per group; scale bar, 100 μm. (B) CNV size on flat-mounts on day 7 in WT mice intravitreally injected with DMSO vehicle or GSK’872, an inhibitor of RIP3 kinase activity. n = 12 eyes per group; scale bar, 100 μm. (C) Western blot analysis of pro-caspase-3 and cleaved caspase-3 in RPE-choroid on day 5 after CNV induction and intravitreal injections with DMSO vehicle, Nec-1 or Nec-1+Z-VAD. n = 3 samples per group. (D) Western blots of caspase 8-containing ripoptosome-like complex immunoprecipitated from RPE-choroid 4 days after CNV induction and intravitreal injections with DMSO vehicle or Nec-1. Both groups eually received Z-VAD intravitreal injections to block the consumption of the formed ripoptosome-like complex. (E) CNV size on flat-mounts on day 7 in WT mice intravitreally injected with DMSO vehicle, Nec-1, or Nec-1+Z-VAD. n = 8 eyes per group; scale bar, 100 μm. (F) Fluorescein leakage from CNV lesions in WT mice was assessed on day 7 and 14 after CNV induction and intravitreal injections with DMSO vehicle, Nec-1 or Nec-1+Z-VAD. n = 30 lesions per group. (G) CNV thickness was assessed on cross-sectional images of CNV obtained through SD-OCT in WT mice on day 7 after CNV induction and intravitreal injections with DMSO vehicle, Nec-1 or Nec-1+Z-VAD. n = 14 lesions per group; scale bar, 100 μm. *P < 0.05, **P < 0.01, ns, not significant; Student’s t-test or one-way ANOVA and post-hoc Tukey’s test. Data are mean ± SEM.

Previous studies have shown that catalytic inhibition of RIP3 leads to caspase activation through kinase-independent function of RIP1/RIP3 through the ripoptosome-like complex comprising RIP1, RIP3, caspase-8 and FADD.(28,29) It also has been shown that RIP1 kinase inhibition by Nec-1 treatment can induce caspase activation and apoptosis.(30,31) Consistent with these studies, RIP1 kinase inhibition leads to caspase-3 activation in CNV lesions, which was abrogated by a caspase inhibitor Z-VAD-FMK (Z-VAD) (Fig. 3C). In contrast, Nec-1 and/or Z-VAD treatment did not alter caspase-3 activation in eyes without CNV lesions (See SI Appendix, Fig. S3). In addition, we directly assessed the formation of ripoptosome-like complex in RPE-choroid tissues with CNV lesions. We observed that Nec-1 treatment upregulated the association of RIP1, RIP3, and FADD with caspase-8, which was clarified by blocking downstream caspase activation that consumes existing ripoptosome-like complex (Fig. 3D).

This activation of caspases by RIP1 kinase inhibition appears to be important for CNV suppression. Indeed, caspase inhibition by Z-VAD abolished the inhibitory effect on CNV by Nec-1 (Fig. 3E). We studied further the effects of Nec-1 and Z-VAD on CNV by employing fluorescein angiography (FA) to evaluate CNV leakage, and spectral-domain optical coherent tomography (SD-OCT) to evaluate CNV size/thickness, as more clinically relevant modalities of CNV assessment. Fluorescein angiography revealed that the severity of vascular leakage from CNV lesions decreased significantly with Nec-1 administration, and that effect was abolished by the pan caspase inhibitor Z-VAD (Fig. 3F), indicating that Nec-1 induced-suppression of angiogenesis is mediated through caspase activation. Similarly, CNV thickness *in situ* was also decreased by Nec-1, an effect reversed through simultaneous injection with Z-VAD (Fig. 3G). Based on all these results, we infer that catalytic inhibition of RIP1 activates caspases to attenuate CNV development.

### RIP1 kinase inhibition suppresses angiogenesis through caspase activation in multiple models *in vivo*

To address whether RIP1 kinase inhibition effects on angiogenesis can be reproduced in other models of angiogenesis, we employed an *in vivo* Matrigel plug assay. Using basic fibroblast growth factor (bFGF), commonly used to induce angiogenesis in Matrigel plugs, we evaluated RIP1 expression in the plugs via immunohistochemistry. We observed higher levels of RIP1 protein in F4/80(+) macrophages, whereas relatively low levels of RIP1 protein in CD31(+) vascular endothelial cells (See SI Appendix, Fig. S4A). Administration of Nec-1 in the Matrigel, decreased bFGF induced angiogenesis as assessed by color (See SI Appendix, Fig. S4B), hematoxylin and eosin (HE) staining (See SI Appendix, Fig. S4C), or hemoglobin content (See SI Appendix, Fig. S4D). Angiogenesis suppression by Nec-1 was abolished by pan-caspase inhibitor Z-VAD (See SI Appendix, Fig. S4B–4D). In addition to bFGF-induced angiogenesis, we also examined the effect of RIP1 kinase inhibition on TNF-related apoptosis-inducing factor (TRAIL)-induced angiogenesis.(32) Again, angiogenesis was significantly suppressed by Nec-1, an effect that was abolished by Z-VAD (See SI Appendix, Fig. S4E–4G).

As a clinically relevant ocular angiogenesis model, alkali injury-induced corneal neovascularization model was used. Alkali injury is an ocular emergency that can lead to severe ocular surface damages. Despite immediate irrigation followed by medical/surgical interventions, a significant number of patients suffer from deteriorated visual function due to the loss of corneal transparency through neovascularization. In this model, we confirmed that RIP1 kinase inhibition by subconjunctival injections of Nec-1 ameliorated corneal neovascularization evaluated 10 days after injury, and the therapeutic effect was abrogated by concomitant Z-VAD injections with Nec-1 (See SI Appendix, Fig. 5).

Taken together, these results suggest that RIP1 kinase inhibition by Nec-1 suppresses angiogenesis through caspase activation in multiple models of angiogenesis.

### Infiltrating macrophages are main target for catalytic inhibition of RIP1 to suppress angiogenesis

Since our data suggested that RIP1 kinase inhibition led to caspase activation, we sought to identify if there is an increase in TUNEL(+) cells and if these cells are macrophages or other cell types. In treatment-naïve CNV lesions, TUNEL(+) cells were most abundantly detected during days 4–5 (See SI Appendix, Fig. S6), the phase of ongoing angiogenic response with abundant macrophage infiltration.(27) Nec-1 administration led to increase in TUNEL(+) cells and this effect was abolished by Z-VAD (Fig. 4A). Consistently, increased number of TUNEL(+) cells were also observed in CNV lesions of RIP1 kinase-inactive RIP1^K45A/K45A^ mice (See SI Appendix, Fig. S7A).(33) Co-immunolabelling to identify these TUNEL(+) cells revealed that they stained positive for the macrophages/microglia marker Iba1 (in cytoplasm) (See SI Appendix, Fig. S7B), but not for endothelial cell marker CD31 (See SI Appendix, Fig. S7C). Based on these results we speculated that Nec-1 treatment induced caspase activation specifically in macrophages. Indeed, on immunohistochemistry cleaved caspase-3 was observed dominantly in F4/80(+) macrophages in the CNV lesions after Nec-1 treatment (Fig. 4B). To exclude the possibility that the cleaved caspase-3 might be due to phagocytosis of apoptotic cells, macrophages were depleted using clodronate, which did not lead to a significant increase of cleaved caspase-3 in non-macrophage cells (Fig. 4B), confirming that the immunolabeled cleaved caspase-3 was from macrophages themselves.

**Fig. 4.**
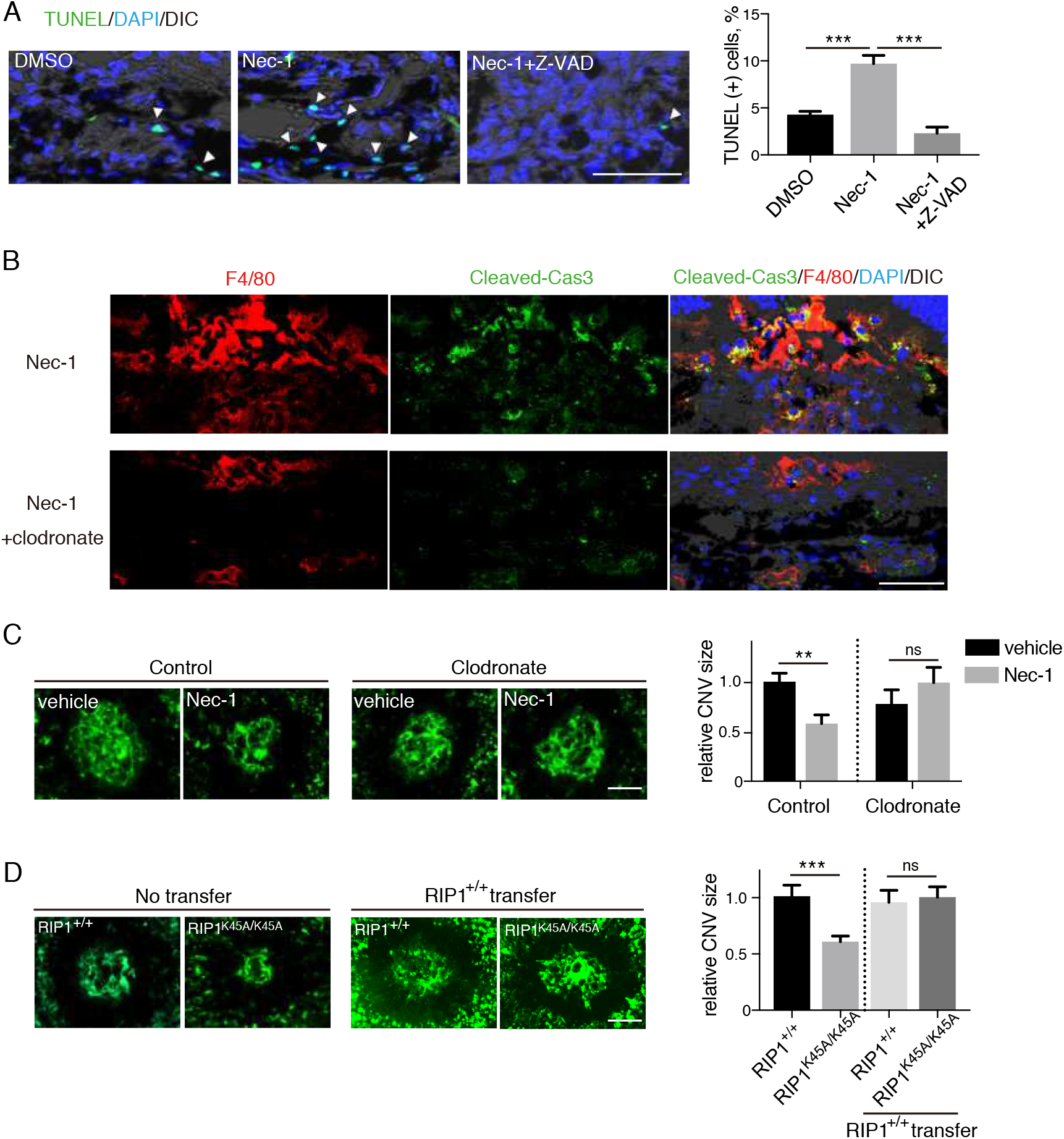
Infiltrating macrophages are targets of catalytic inhibition of RIP1 to attenuate CNV. (A) TUNEL staining of CNV sections on day 5 after CNV induction and intravitreal injections of with DMSO vehicle, Nec-1 or Nec-1+Z-VAD. n = 6 lesions per group; scale bar, 50 μm. (B) Immunohistochemistry of day 4 CNV sections for cleaved caspase-3 and its localization in F4/80(+) macrophages after intravitreal injections (day 0) with Nec-1 and intraperitoneal injections (day 2) with vehicle or clodronate. Scale bar, 50 μm. (C) CNV was induced in WT mice and DMSO vehicle or Nec-1 was intravitreally injected on day 0. The mice received intraperitoneal injections with control or clodronate liposome on day 2. CNV size was assessed on flat-mounts on day 7. n = 8 eyes per group; scale bar, 100 μm. (D) CNV size on flat-mounts on day 7 in RIP1^+/+^ and RIP1^K45A/K45A^ kinase dead mice. The designated groups of mice received an adoptive transfer of bone marrow monocytes (5×10^6^) from RIP1+/+ mice on day 2. n = 10 eyes per group; scale bar, 100 μm. **P < 0.01, ***P < 0.001; Student’s t-test or one-way ANOVA and post-hoc Tukey’s test. ns, not significant difference. Data are mean ± SEM.

The above-mentioned results suggest that infiltrating macrophages may be the targets for the catalytic inhibition of RIP1 to suppress angiogenesis. To further address this hypothesis, WT mice were divided into 2 groups; one group received intraperitoneal injections of clodronate liposomes at 2 days after CNV induction to deplete macrophages, and the other group received intraperitoneal injections of control liposomes. Nec-1 did not suppress CNV development in mice with macrophage depletion, while it did suppress CNV development in non-depleted mice (Fig. 4C). Furthermore, we employed RIP1 kinase-inactive (RIP1^K45A/K45A^) mice, and confirmed that the genetic inhibition of RIP1 kinase activity reduced CNV size (Fig. 4D) as did pharmacological RIP1 kinase inhibition using Nec-1. To clarify the role for RIP1 kinase inhibition in macrophages, bone marrow monocytes of WT mice were adoptively transferred to the intraperitoneal cavity of WT mice or RIP1^K45A/K45A^ mice on day 2 after CNV induction (See SI Appendix, Fig. S8). After the adoptive transfer, the CNV size in RIP1^K45A/K45A^ was significantly increased to the similar size of the WT mice (Fig. 4D). Taken together, these data suggest that the catalytic inhibition of RIP1 suppresses angiogenesis, at least partially, through action on infiltrating macrophages.

### Effects of RIP1 kinase inhibition on macrophage function

Although RIP1 kinase inhibition in macrophages and their effect on angiogenesis could be mediated by increased macrophage death and elimination, it could also be a result of non-cell-death-mediated effects on macrophages’ function. Thus, to examine for this later potential we investigated differences in the expression levels of M1-like and M2-like macrophage markers in CNV lesions. We observed lower mRNA levels of markers for M2-like macrophages (Arg-1, Fizz1, Ym1/2, MMP9, MMP12, IL-10, PDGF-B, Wnt5a, Wnt7b, IL-4, and IL-13) in kinase-dead RIP1^K45A/K45A^ compared to WT and no significant difference in the mRNA levels of markers for M1-like macrophages (iNOS, TNF, IL-6, IL-18, IL-23a, Cxcl1, Cxcl2, Cxcl10, and Cxcl11) (Fig. 5A). We also assessed protein levels of VEGF and IL-12 using ELISA. VEGF-A is a marker for pro-angiogenic M2-like macrophages, whereas IL-12 is a marker of M1-like macrophages and a well-established anti-angiogenic cytokine.(34–36) We found that VEGF-A protein levels were down-regulated, whereas IL-12 protein levels were up-regulated in eyes with CNV lesions from RIP1^K45A/K45A^ mice compared with WT mice (Fig. 5B). These data suggest that RIP1 kinase inhibition may not only upregulate caspase-mediated cell death in macrophages but may also modulate M1/M2 polarization. Consistent with this impression, phosphorylation and activation of STAT1, a key transcription factor for M1 macrophages, was upregulated in CNV lesions of RIP1^K45A/K45A^ compared to WT, whereas phosphorylation and activation of STAT6, a key transcription factor for M2 macrophages was suppressed (Fig. 5C). In addition, immunohistochemistry revealed down-regulated M2 marker CD206 expression in the CNV macrophages of RIP1^K45A/K45A^ mice (Fig. 5D).

**Fig. 5.**
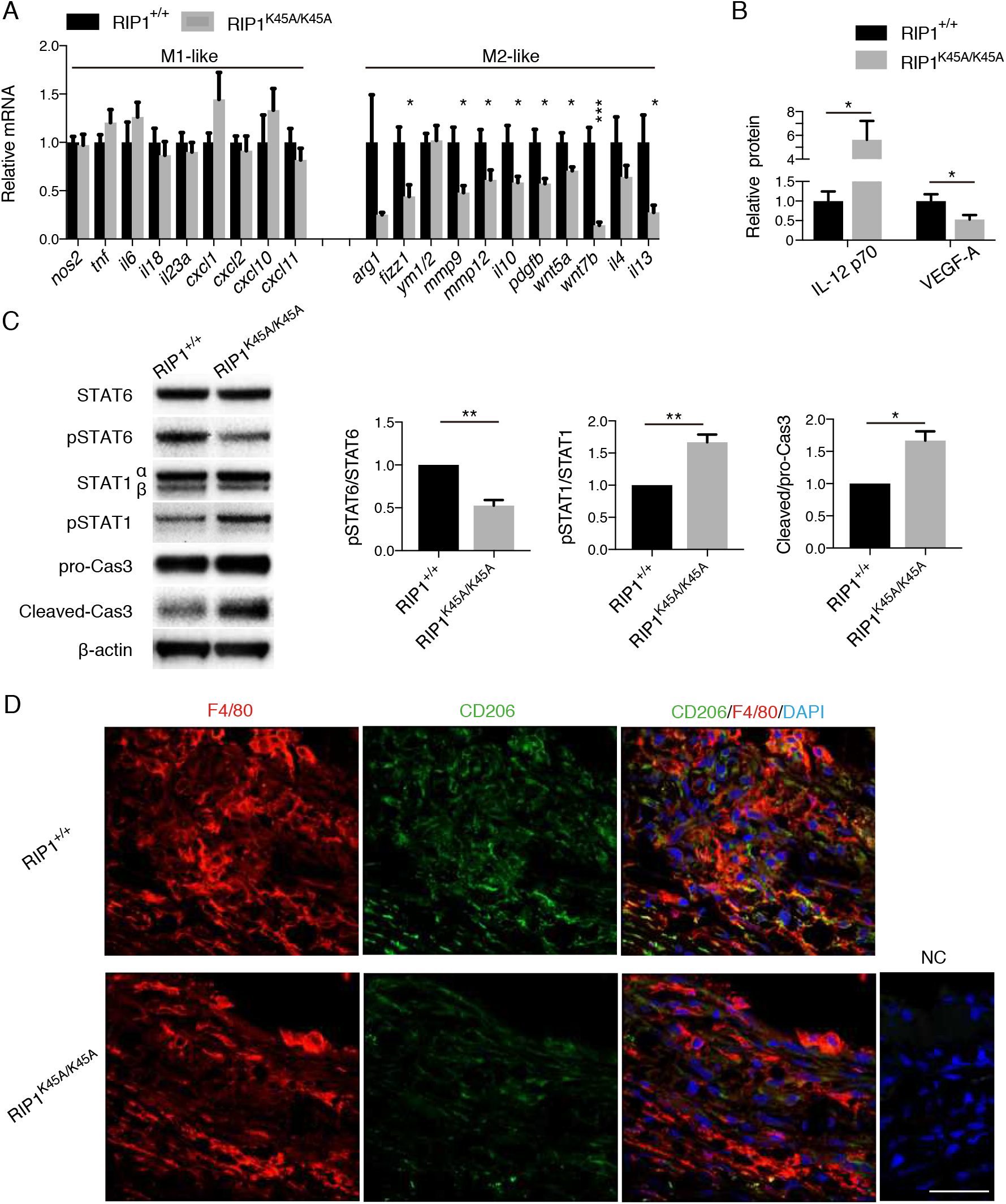
(A) Relative mRNA levels on day 4 after CNV induction in RPE-choroid samples of RIP1+/+ and RIP1K45A/K45A mice. n = 6 eyes per group. (B) Relative protein levels on day 5 after CNV induction in RPE-choroid samples of WT and RIP1^K45A/K45A^ mice. n = 6 eyes per group for IL-12, n = 14 per group for VEGF-A. (C) Western blot analysis of STAT6, phospro-STAT6 (Tyr641), STAT1, phosphor-STAT1 (Tyr701), pro-caspase-3 and cleaved caspase-3 in RPE-choroid on day 4 after CNV induction in RIP1^+/+^ and RIP1^K45A/K45A^ mice. n = 3 samples per group. (D) Immunohistochemistry of day 4 CNV sections for CD206 and its localization in F4/80(+) macrophages in RIP1+/+ and RIP1^K45A/K45A^ mice. Scale bar, 50 μm. *P < 0.05, **P < 0.01, ***P < 0.001, ns, not significant; Student’s t-test or one-way ANOVA and post-hoc Tukey’s test. Data are mean ± SEM.

### Inhibition of RIP1 kinase activity suppresses M2 polarization of macrophages *in vitro*

The *in vivo* results suggested that catalytic inhibition of RIP1 has an additional non-apoptotic function to modulate macrophage activation altering M1/M2 polarization. To further evaluate this, we used bone marrow-derived macrophages (BMDMs) *in vitro* and used IL-4 to induce M2 phenotype. After 24 h, RIP1 kinase inhibitor Nec-1 was added to the culture medium to levels not affecting viability ((37,38) and See SI Appendix, Fig. S9) in order to assess cell death-independent roles of RIP1 in M2 macrophages. Inhibition of RIP1 kinase by Nec-1 in IL-4 treated BMDMs resulted in caspase-8 activation and suppression of the critical M2 transcription factor STAT6 (Fig 6A). Both effects were reversed by pan-caspase inhibitior Z-VAD (Fig. 6A). Analysis by flow cytometry of M2 (IL-4) markers CD206(+) or CD163(+) showed a significant reduction after RIP1 kinase inhibition, which was also rescued by Z-VAD (Fig. 6B). Similarly, production of M2 cytokine IL-10 was suppressed with RIP1 kinase inhibition and was rescued by Z-VAD (Fig. 6C). Finally, we assessed mRNA levels of macrophage polarization. After a 24 h treatment with Nec-1, the mRNA levels of M2 markers (Arg-1, Fizz1, and Ym1/2) were significantly down-regulated, and this effect was reversed by Z-VAD (Fig. 6D). With extended treatment with Nec-1 for 48 h, we observed significant up-regulation of mRNA levels of M1 markers (iNOS, IL-6, IL-23a, Cxcl1, Cxcl2, il12a, and Cxcl11), which was also abolished by Z-VAD treatment (Fig. 6D). These data support that the catalytic inhibition of RIP1 suppresses M2 polarization and may shift macrophages to M1 type polarization at least partially through the activation of the caspase pathway. In addition, to rule out the off-target effects of Nec-1 to modulate macrophage polarization, BMDMs from RIP1 kinase-dead RIP1^K45A/K45A^ mice were used to confirm that genetic RIP1 kinase inhibition also suppressed M2 marker expressions while enhanced M1 marker expressions on mRNA levels (See SI Appendix, Fig. S10).

**Fig. 6.**
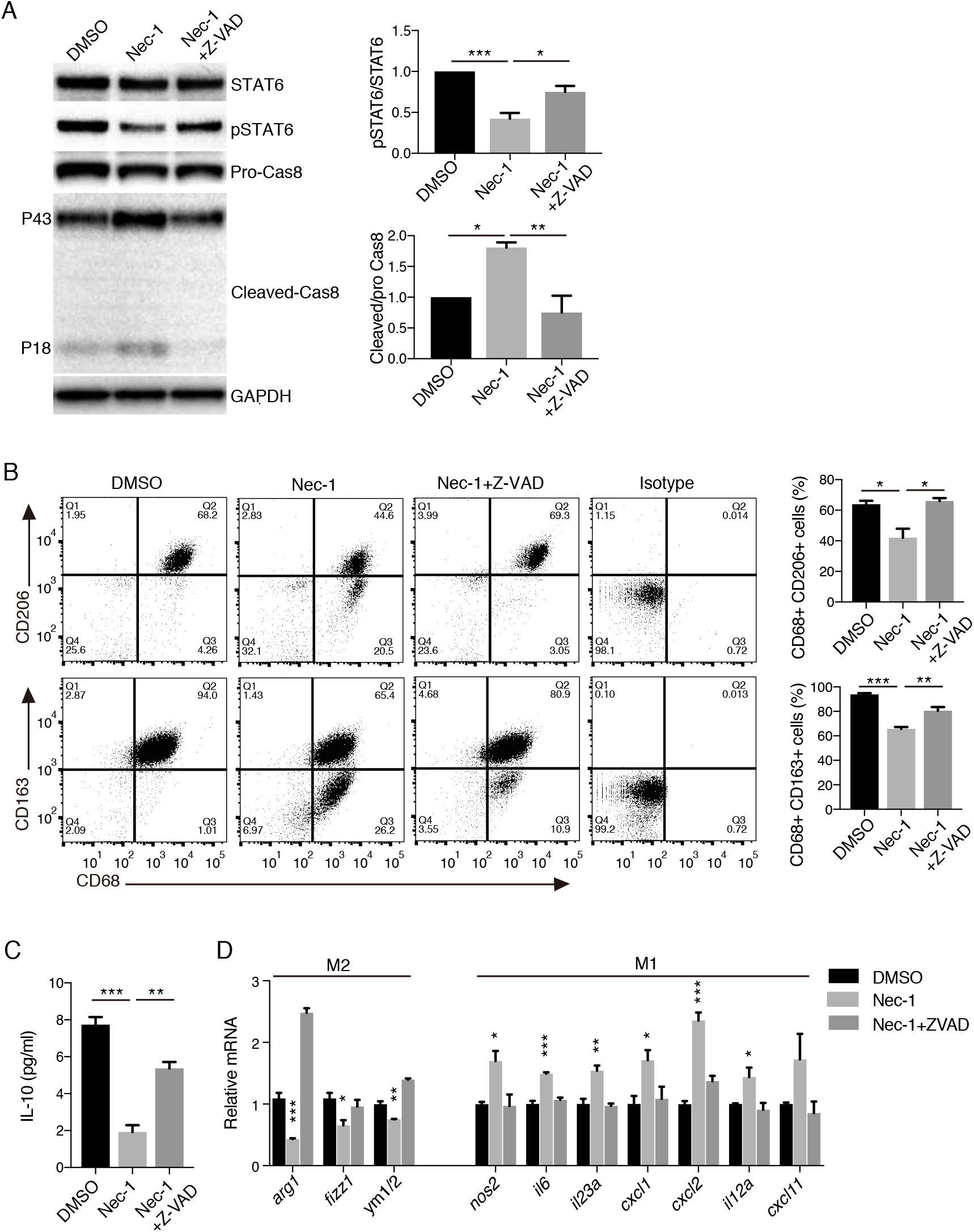
Nec-1 suppresses M2 polarization of BMDM in vitro through caspase activation. (A) Western blot analysis of STAT6, phospho-STAT6 (Tyr641), pro- caspase-8, and cleaved caspase-8 in M2-polarized BMDM by IL-4 and treated with DMSO vehicle, Nec-1 (30 μM), or Nec-1 (30 μM) + Z-VAD (30 μM) for 48 h. n = 3 samples per group. (B) Flow cytometry analysis of M2-polarized BMDM treated with Nec-1 (30 μM), or Nec-1 (30 μM) + Z-VAD (30 μM) for 48 h. Cells were stained for CD68, CD206, and CD163. n = 3 samples per group (C) IL-10 secretion in supernatant by M2-polarized BMDM treated with Nec-1 (30 μM), or Nec-1 (30 μM) + Z-VAD (30 μM) for 48 h. n = 5 samples per group (D) Relative mRNA levels of M2-polarized BMDM treated with Nec-1 (30 μM), or Nec-1 (30 μM) + Z-VAD (30 μM) for 24 h (for M2 markers) and 48 h (for M1 markers). n = 3–4 samples per group. *P < 0.05, **P < 0.01, ***P < 0.001; one-way ANOVA and post-hoc Tukey’s (A–C) or Dunnet’s test (D). Data are mean ± SEM.

### Catalytic inhibition of RIP1 does not affect the angiogenic response in endothelial cells

RIP1 expression is considered ubiquitous, and our histological analyses had shown that in addition to infiltrating macrophages RIP1 expression was also seen at lower levels in vascular endothelial cells, suggesting that they may also play a role in mediating the effects of RIP kinase inhibition on angiogenesis. To assess this possibility, we examined the effects of Nec-1 treatment *in vitro* using cultured human umbilical vein endothelial cells (HUVECs). RIP1 kinase inhibition did not decrease proliferation of HUVECs for up to 87 h of culture compared to vehicle (See SI Appendix, Fig. S11A). Additionally, RIP1 kinase inhibition did not affect migration of HUVECs tested by the scratch-wound assay (See SI Appendix, Fig. S11B), nor tube formation assay (See SI Appendix, Fig. S11C). Furthermore, we evaluated *ex vivo* choroidal angiogenesis by culturing 1 x 1 mm pieces of peripheral RPE-choroid-sclera on Matrigel as described previously.(39–41) This system enables the assessment of choroidal angiogenesis without infiltrating macrophages. Consistent with the results of HUVECs, Nec-1 treatment did not suppress choroidal angiogenesis *ex vivo* (See SI Appendix, Fig. S11D). These results taken together suggest that endothelial cells are not the direct targets for the catalytic inhibition of RIP1 to attenuate angiogenesis.

## DISCUSSION

RIP kinases have been extensively studied as the key effectors of regulated cell death (necroptosis) and their role has been shown in many disorders including neurodegenerative diseases,(42–45) ischemic injuries,(46) inflammatory diseases,(47) and cancer metastasis.(48) However, the previous studies have not clarified the role of RIP kinases in vascular disorders. The present study demonstrates for the first time a non-necrotic role of RIP1 kinase activity in angiogenesis. We showed that the pharmacological and genetic inhibition of RIP1 kinase activity is effective in inhibiting angiogenesis in multiple *in vivo* models. Mechanistically, this effect was not mediated by RIP kinase inhibition in endothelial cells, but appeared to be mediated at least partially through RIP kinase inhibition-mediated caspase activation through ripoptosome-like complex and suppression of M2 infiltrating macrophages. In contrast, caspase inhibition, which leads to activation of necroptotic pathway,(19–21) aggravated CNV *in vivo* (See SI Appendix, Fig. S12A), and enhance M2 marker expressions in BMDMs *in vitro* (See SI Appendix, Fig. 12B). A schematic drawing (Fig. 7) shows the mechanism that the balance between apoptotic and necroptotic pathways in macrophages may dictate pathological angiogenesis.

**Fig. 7.**
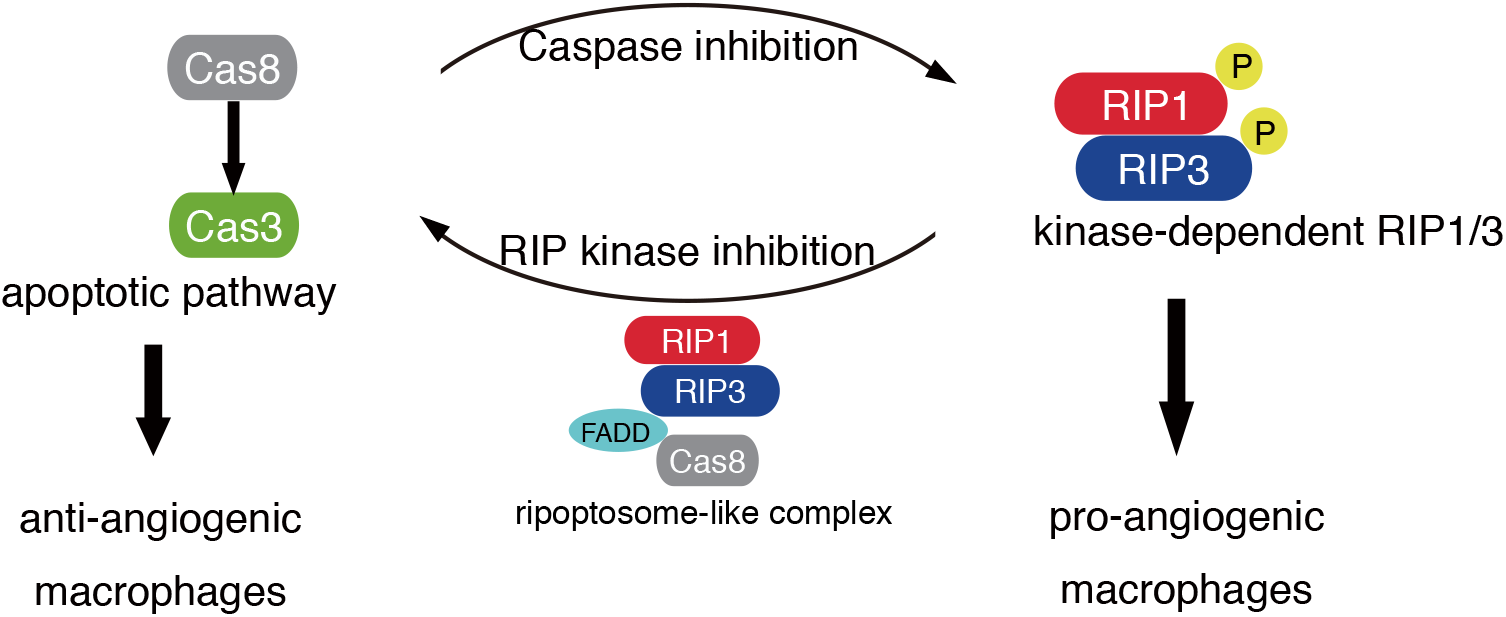
Schematic summary for the present study. In angiogenesis-associated macrophages, Nec-1 treatment or kinase-inactive RIP1 activates caspases through kinase-independent ripoptosome-like complex and suppress M2 activation, thereby inhibits angiogenesis. In contrast, caspase inhibition that augments RIP kinase activation aggravates pathological angiogenesis.

Recent studies have unveiled new levels of complexity in cell fate regulation by RIP1. For example, while RIP1 can activate RIP3, it also acts as a negative regulator of spontaneous RIP3 activation and necroptosis.(23) Other experiments have revealed an essential function of RIP1 in preventing inappropriate activation of both caspase-8-dependent apoptosis and RIP3-dependent necroptosis in multiple tissues, including skin, liver, and intestine.(24–26) In addition, RIP1 kinase-dependent apoptosis has also been observed in TNF-induced cell death, in which phosphorylation at the intermediate domain of RIP1 (Ser321 for mouse and Ser320 for human RIP1) controls RIP1 kinase-dependent apoptosis and necroptosis.(49,50) Overall, accumulating data have revealed multiple roles for RIP1 in determining cellular fate, depending on cell types and contexts: nevertheless, a complete understanding remains elusive. Similar to RIP1, the role of RIP3 varies depending on the biological milieu. In addition to the necroptotic pathway induced by kinase-dependent RIP3 activity, a recent study revealed that genetic (RIP3^D161N/D161N^) or pharmacological inhibition of RIP3 kinase inhibition leads to apoptosis via assembly of ripoptosome-like complex harboring kinase-independent RIP1/RIP3, FADD, and caspase-8.(28,29) These findings as well as the previous reports of Nec-1-induced apoptosis(30,31) may suggest the reciprocal regulation between apoptosis and necroptosis, in which caspase-8 blocks necroptosis through RIP1 cleavage while RIP1/RIP3 kinase activity blocks apoptosis through unknown mechanism.(51) Kinase-inactive RIP3^D161N/D161N^ knock-in mice die during embryogenesis. Kinase-inactive RIP3^K51A/K51A^ mice are viable, but display greatly reduced expression of RIP3.(20) In addition, kinase-independent RIP3 function has also been proposed to control cytokine production in mononuclear phagocytes in the lamina propria of the intestine after treatment with dextran sodium sulfate.(52)

In the present study, RIP up-regulation was observed in lesions with new vessel formation and RIP kinase inhibition suppressed angiogenesis formation. Because RIP1 is a ubiquitously expressed protein, we sought to examine which cell type was important for the observed phenomenon. Experiments in isolated cultured endothelial cells and in vascular explants without immune cells showed that RIP1 kinase inhibition did not affect vascular endothelial cell proliferation, tube formation or vascular sprouting (See SI Appendix, Fig. S11), suggesting that the important cell type in RIP1 kinase and angiogenesis may be the immune cells and not the endothelial cell. Thus, we next turned our attention to the immune cells. First, when macrophages were depleted using clodronate, Nec-1 could not suppress neovascularization. Second, adoptive transfer of WT macrophages to RIP1^K45A/K45A^ mice dampened the ameliorating effect of RIP1 ^K45A/K45A^ on neovascularization. These results suggest that the macrophages play the important role of RIP1 kinase in angiogenesis. But how was RIP1 kinase inhibition suppressing angiogenesis? Because RIP kinases are essential in regulated cell death called necroptosis, we first examined that proposition. However, although the observed angiogenesis was suppressed through the catalytic inhibition of RIP1 or RIP3 (Nec-1, GSK’872, or kinase-dead RIP1^K45A/K45A^), it was not suppressed after total deletion of RIP3, suggesting that the inhibitory effect on angiogenesis does not reflect the inhibition of the necroptotic pathway alone.

Previous reports have suggested that kinase inhibition of RIP3 unmasks a kinase-independent function of RIP3 that leads to activation of caspase-8.(28,29) Similarly studies have reported caspase-dependent apoptosis induced by Nec-1 treatment and inhibition of RIP1 kinase.(30,31) Based on this knowledge we speculated that caspase activation by kinase-independent RIP function (which was unmasked by RIP kinase inhibition(28,29)) contributes to the attenuation of angiogenesis. Indeed, we observed increased caspase activation in macrophages in CNV of Nec-1-treated eyes, and caspase inhibition by Z-VAD clearly blocked the inhibitory effect of Nec-1 on angiogenesis in laser-induced CNV, Matrigel plugs, and alkali injury-induced corneal neovascularization in mice. Furthermore, we observed an increased number of TUNEL(+) macrophages and caspase-3 activation by RIP1 kinase inhibition in CNV, and these effects were blocked by Z-VAD, suggesting that the apoptotic function of caspase was induced by RIP1 kinase inhibition in macrophages. Although induction of apoptotic cell death in macrophages could partially explain the suppression of angiogenesis (as is known that macrophage depletion reduces angiogenesis(53,54)), the number of TUNEL(+) cells in CNV lesions was relatively small, and thus apoptotic function of caspase by itself in macrophages may not be able to totally explain the angiogenesis suppression seen. Thus, we looked for other non-cell-death effects of RIP1 kinase inhibition and caspase activation in macrophage functions. Since M2 type of macrophages are considered the pro-angiogenic type of macrophages we investigated if RIP1 kinase inhibition and caspase activity can alter M1/M2 polarization. Indeed, after RIP1 kinase inhibition we observed down-regulated M2-like marker expressions and up-regulated M1-like marker expressions in macrophages treated with Nec-1 and the effect was reversed by Z-VAD, suggesting non-apoptotic function of caspase by RIP1 kinase inhibition to suppress angiogenic phenotypes of macrophages *in vivo*. This was further corroborated by *in vitro* studies with BMDMs.

Previous work has shown that caspase-8 is not only an initiator of apoptotic cell death but also plays a non-apoptotic role in facilitating monocyte-macrophage differentiation.(55–57). In addition, a previous study has shown that a caspase inhibitor IETD-CHO promotes M2 macrophage polarization in vitro, which is in line with our hypothesis (Fig. 7).(58) Another recent study also reported a vital role of caspase-8 for M1 polarization. In caspase-8 deficient macrophages RIP1/RIP3 necroptotic pathway was hyper-activated, which prevented the normal M1 polarization.(59) These studies have revealed a cell death-independent role of caspase for regulating M1 macrophage polarization. In contrast to these previous studies revealing a role for caspase for M1 polarization, we elucidate in the current study that RIP1 governs macrophage polarization by facilitating M2 polarization through cell death-independent functions. A very recent study has just reported the role of RIP1 to facilitate M2 polarization in tumor-associated macrophages,(60) which is consistent with our conclusion. We further revealed the mechanistic aspects in which RIP1 kinase inhibition unmasked the kinase-independent function of RIPs, leading to formation of ripoptosome-like complex formation and caspase activation. Hence, our current study suggests a novel association between caspase and necroptotic pathways to determine M1/M2 macrophage polarization.

We enrolled two ocular angiogenesis models (CNV and alkali injury) and Matrigel plug model, and revealed the similar necroptotic and apoptotic roles in multiple *in vivo* models. However, different mechanisms may govern in different angiogenesis models, and further studies would be necessary for other angiogenesis models. For example, some researchers have suggested naturally occurring retino-choroidal anastomotic neovascularization in some mouse strains might be useful for human AMD (61), although the phenotype mimics a relatively small portion of human AMD.

In summary, we have identified a previously unsuspected role for RIP1 kinase activity in angiogenesis mediated at lest partially through action on macrophages. These results provide evidence for the therapeutic value of the catalytic inhibition of RIP1 to attenuate angiogenesis, which depends on macrophage-specific caspase activation. These findings are expected to shed new light on the pathophysiology where the balance between RIP1 kinase activity and caspase activity is described to play an important role in macrophage activation for angiogenesis.

## MATERIAL AND METHODS

Animal procedures were approved by the Animal Care Committee of Massachusetts Eye and Ear, and performed in accordance with the Association for Research in Vision and Ophthalmology Statement for the Use of Animals in Ophthalmic and Vision Research. RIP1^K45A/K45A^ mice were kindly provided by GlaxoSmithKline (Brentford, Middlesex, UK). Rip3^−/−^ mice and C57BL/6J mice were purchased from Jackson Laboratory (Bar Harbor, ME, USA). These animals were backcrossed to C57BL/6J mice to remove the Crb1^rd8/rd8^ mutation.

Detailed descriptions on laser-induced CNV model, immunohistochemistry, western blot, *in vivo* Matrigel plug assay, TUNEL staining, quantitative real-time RT-PCR, ELISA, HUVEC assays, ex vivo choroidal sprouting assay, BMDM culture, and flow cytometry can be found in *SI Materials and Methods*.

## Acknowledgements

This work was supported by the Yeatts Family Foundation (D.G.V.); 2013 Macula Society Research Grant award (D.G.V.); a Physician Scientist Award (D.G.V.); unrestricted grant from the Research to Prevent Blindness Foundation (J.W.M. and D.G.V); NEI R21EY023079-01/A1 (D.G.V.); NEI grant EY014104 (MEEI Core Grant) (D.G.V.); Loeffler Family fund (D.G.V.); R01EY025362-01 (D.G.V.); ARI Young investigator Award (D.G.V.); Foundation Lions Eye Research Fund (D.G.V.); NIH NEI Core grant P30EY003790 (D.G.V.); Agence National de la Recherche (ANR), the ERA-Net for Research on Rare Diseases, Institut Universitaire de France (D.G.V.), and fellowship from TOYOBO Biotechnology Research Foundation (T.U.).

## Author Contributions

T.U. and D.G.V. conceived and designed the research. T.U., S.N., J.J.L., K.I., K.A., T.T., and E.P. conducted the research. D.M., N.E. and E.H. provided necessary materials and experimental protocols. T.U., S.N., Y.M. and D.G.V. performed statistical analysis and/or interpreted data. T.U., Y.M. K.I. and D.G.V. wrote the manuscript. D.G.V. supervised the research. J.W.M., D.G.V., and M.A. provided resources for the conduct of the research.

## Competing Financial Interests statement

The authors declare no competing financial interests.

