## Supporting Information for "RIP1 kinase mediates angiogenesis by modulating macrophages in experimental neovascularization"

### SI Materials and Methods

#### Laser-Induced CNV Model

The protocol was performed as previously described (1–4). Eight-week-old mice were anesthetized using tribromoethanol (avertin) solution and the pupils were dilated with 5%-phenylephrine and 0.5%-tropicamide ophthalmic solution. Laser photocoagulation was performed using a 532-nm laser (Oculight GLx Laser System, IRIDEX Corporation, Mountain View, CA, USA) attached to a slit lamp. A cover slip was used as a contact lens through which the mouse fundus was observed. Laser spots were generated using a green laser (532 nm wavelength, 100 µm spot size, 100 msec duration, and 100 mW power). The laser spots were located two to three disc diameters from the optic nerve head. Successful laser photocoagulation was confirmed by immediate bubble formation resulting from the rupture of Bruch's membrane. Four laser spots were generated per eye to evaluate CNV size, fluorescein angiography, optical coherence tomography (OCT), and immunohistochemistry. In contrast, 40 laser spots were generated per eye to evaluate mRNA and protein expressions. For the designated experiments, 0.2 ml of clodronate liposome or control liposome (Encapsula NanoSciences LLC, Brentwood, TN, USA) was intraperitoneally administered.

Intravitreal injections of Nec-1 was performed using 1.5µl DMSO vehicle at a concentration of 100 µM (lower) or 300 µM (standard). Z-VAD and GSK'872 were intravitreally injected as 1.5µl DMSO solution at a concentration of 300 µM and 100 µM, respectively. The concentration is diluted 10 times in the intraocular space. Nec-1 (15 mg/kg/day) or vehicle (DMSO) was also administered through subcutaneous osmotic pumps (Model 1002, Alzet, Cupertino, CA, USA) implanted the day before CNV induction. Because CNV size can be affected by the intravitreal injections of DMSO vehicle depending on the dose and the timing after CNV induction, relative CNV size was demonstrated compared to the control group of each experiment.

On day 7 or 14, the mice were anesthetized and perfused through the left ventricle with 1ml of fluorescein concanavalin A (FL-1001; Vector Laboratory, Burlingame, CA, USA) diluted ten times with PBS. The mice were sacrificed, and the eyes were subsequently enucleated and fixed in 4% paraformaldehyde for 20 min, followed by PBS replacement. Under the microscope, eyecups were generated and the RPE-choroid-sclera was flat mounted onto slides using mounting medium and coverslips. Images of CNV were captured using a fluorescein microscope (Zeiss Axio Imager.M2, Thornwood, NY, USA). Blinded examiners measured the area of each CNV lesion using Photoshop CS3 software.

On day 7, cross-sectional images of CNV *in situ* were obtained using an SD-OCT system (Bioptigen Inc., Durham, NC, USA). Each scan area was 1.4 × 1.4 mm that covers a whole CNV lesion. Each area was scanned with 100 horizontal, raster, and consecutive B-scans. Each B-scan consists of 1,000 A scans. The thickness of each CNV lesion, defined as the distance from the outer boundary of the RPE layer to the inner tip of the CNV lesion, was evaluated.

On day 7 and 14, fluorescein angiography was performed using Micron III (Phoenix Research Labs, Pleasanton, CA, USA). The images of fluorescein angiogram were evaluated based on the established grading scales (5,6); bright hyperfluorescence in the transit images and late leakage beyond the treated areas (grade IV) were considered clinically significant.

For the designated laser-induced CNV experiments, adoptive transfer of bone marrow monocytes was performed. For this purpose, bone marrow cells were isolated and cultured in the presence of M-CSF as described below (BMDM culture section) for 6 days. The adherent cells were collected using a scraper, and 5 x $10^6$ monocytes were intraperitoneally injected to the mice after 2 days of CNV induction. The procedures were validated using monocytes of EGFP mice that infiltrate to the CNV lesions on day 4 (Fig. S7)

**Immunohistochemistry**
After euthanization, the enucleated mouse eyes or Matrigel plugs were fixed in 4% paraformaldehyde for 2 or 4 h, respectively, at room temperature (RT), dehydrated in 30% sucrose at 4°C overnight, and embedded in OCT compound. The eyes and Matrigel samples were cryosectioned at 10- and 16-μm thicknesses, respectively. For immunofluorescence, the sections were permeabilized using -20°C methanol for 5 min, and subsequently, the slides were treated with antigen retrieval solution (HistoVT One, Nacalai Tesque, Kyoto, Japan) at 70°C for 20 min. After blocking with PBS-TB (0.3% Triton and 0.5% BSA) for 1 h at RT, the slides were incubated with primary antibodies (Table S1) overnight at 4°C. Subsequently, the slides were washed with PBS, incubated with secondary antibodies (Thermo Fisher Scientific, Waltham, MA, USA) for 1 h at RT, washed with PBS, and coverslipped using PloLong Gold Antifade Reagent (Thermo Fisher Scientific). Images were captured using a confocal microscope (LSM800, Zeiss, Oberkochen, Germany).
For paraffin sections of human eyes with neovascular AMD, The deidentified eye was obtained through National Disease Research Interchange (Philadelphia, PA). The sections were deparaffinized and immunostained using ImmPRESS Excel Staining Kit (Vector Laboratories) according to the manufacturer’s instructions. Sections were conterstained with hematoxylin.

**Western Blot and immunoprecipitation**
Tissues or cultured cells were collected in RIPA buffer, homogenized, and sonicated briefly. The samples were heated in NuPAGE Sample Buffer (Thermo Fisher Scientific) containing 2-mercaptoethanol at 95°C for 5 min and electrophoresed using NuPAGE Bis-Tris Gels (Thermo Fisher Scientific) at 150 V.

The proteins in the gels were transferred to PVDF membranes using a semi-dry protocol. The membranes were blocked with 5% skim milk in PBS-T (0.1% Tween 20) at RT for 1 h, washed, incubated with primary antibodies (Table S1) in the blocking solution overnight at 4°C, washed, incubated with HRP-conjugated secondary antibodies in blocking solution for 1 h, and thoroughly washed. The blots were imaged using Amersham ECL Select Western Blotting Detection Reagent (GE Healthcare Life Sciences, Chicago, IL, USA) and the ChemiDoc imaging system (Bio-Rad, Hercules, CA, USA). For immunoprecipitation, lysate samples were incubated with rat anti-caspase-8 antibody (Enzo Life Sciences) overnight at 4°C, pulled down using protein G-agarose beads (Cell Signaling Technology), and eluted by heating in SDS sample buffer. Immunoprecipitated samples were blotted using mouse anti-RIP1 (BD Biosciences), rabbit anti-RIP3 (NOVUS), rabbit anti-caspase-8 (Cell Signaling Technology), and mouse anti-FADD (Santa Cruz) antibodies. Specificity of the caspase-8 antibody used for immuneoprecipitation was confirmed (See SI Appendix, Fig. S13).

#### *In Vivo* Matrigel Plug Assay

Experiment was performed as previously described (7,8). A mixture of bFGF (final concentration 150 ng/ml) and 20 units of heparin, or TRAIL (final concentration 400 ng/ml) was dissolved in Growth Factor Reduced Matrigel (Corning Inc., Corning, NY, USA) to induce angiogenesis in the Matrigel plugs. For treatment, vehicle DMSO, Nec-1 (30 μM), or Nec-1 (30 μM) with Z-VAD (30 μM) was dissolved in the Matrigel. Mice were anesthetized, and 200 μl of Matrigel was injected at the ventral surface in the groin area close to the dorsal midline. On day 7 after the injections, the mice were euthanized, and the Matrigel plugs were excised. Each plug was homogenized in 500 μl of PBS containing 20 units of heparin, and the hemoglobin content was quantified using a colorimetric assay kit (Cayman Chemical, Ann Arbor, MI, USA). The value was adjusted by the weight of each plug.

#### Alkali injury-induced corneal neovascularization

Alkali injury was induced by 2 mm-diameter filter paper soaked with 1.0 M sodium hydroxide (NaOH) that was applied on the mouse cornea for 30 seconds. Then the eyes were immediately irrigated with 20 ml PBS (Day 0). From Day 3–9, the injured eyes were treated with subconjunctival injections (once daily) of 20μl DMSO vehicle, Nec-1 (300 μM), or Nec-1 (300 μM) with Z-VAD (300 μM) solutions. On day 10, the mice were anesthetized and perfused with fluorescein concanavalin A (FL-1001; Vector Laboratory). The mice were sacrificed, and the eyes were enucleated and fixed in 4% paraformaldehyde, followed by PBS replacement. Under the microscope, the cornea was isolated and flat mounted onto slides using mounting medium and coverslips. Images were captured using a fluorescein microscope. Blinded examiners measured the area of each CNV lesion using Photoshop CS3 software.

#### TUNEL staining

DNA breaks in CNV lesions were detected on the cryosections through the TUNEL assay using the ApopTag Plus In Situ Apoptosis Fluorescein Detection Kit (Thermo Fisher Scientific), which utilizes a fluorescein-conjugated anti-digoxigenin antibody.

**Quantitative Real-Time RT-PCR**

RNA was extracted using TRIzol Reagent (Thermo Fisher Scientific). The cDNA was synthesized using Superscript VILO Master Mix (Thermo Fisher Scientific). The cDNA samples were mixed with Fast SYBR Green Master Mix (Thermo Fisher Scientific) and primer pairs, and real-time RT-PCR was conducted using the QuantStudio 5 Real-Time PCR System (Thermo Fisher Scientific). The primers used in the present study are listed in Table S2. The mRNA levels were quantified as the difference in cycle threshold values (ΔΔCt), and GAPDH was used as the inner control.

**ELISA**

The tested samples were either supernatant of the cultured cells or mouse eyes homogenized in NP-40 lysis buffer. ELISA kits for mouse VEGF-A, IL-12p70, and IL-10 in which monoclonal antibodies were pre-coated onto the microplates were used (all from R&D Systems). These kits employ a quantitative sandwich enzyme immunoassay technique.

**HUVECs**

HUVECs were purchased from Thermo Fisher Scientific. HUVECs were seeded at a concentration of 2.0 x $10^5$ cells/ml and cultured in the medium 200PRF with Low Serum Growth Supplement (Thermo Fisher Scientific). Nec-1 (30 μM), Z-VAD (30 μM), and DMSO vehicles were contained in the culture medium from 0 h (i.e., when the cells were seeded). For the proliferation assay, 100 μl/well of the cell suspension was used for 96-well plate. The dehydrogenase activity in live cells was measured using a CCK-8 (Dojindo, Kumamoto, Japan) assay kit, similar to the MTT assay. For migration assay, HUVECs were grown until confluence in 24-well plates. A scratch wound was generated using a sterile 100-μl tip in each well. The medium was changed, and treatment with Nec-1 (30 μM), Z-VAD (30 μM), and DMSO vehicles was initiated (0 h). Images of the scratch wounds were taken using a phase-contrast microscopy (Primeovert, Zeiss) at 0 and 10 h, and the distance of the wound was measured using Photoshop CS3 software. For the *in vitro* tube formation assay, each well of the 24-well plate was covered with 150 μl of Geltrex (Thermo Fisher Scientific), and the plate was incubated for 30 min at 37°C to allow the gel to solidify. A 400-*μ*l aliquot of cell suspension containing Nec-1 (30 μM), Z-VAD (30 μM), and DMSO vehicles was added to each well and incubated at 37°C with 5% $CO_2$ for 16 h. Images were captured using a phase-contrast microscopy (Primeovert, Zeiss), and the number of mesh/field were counted.

***Ex Vivo* Choroidal Sprouting Assay**

Sprouting of choroidal vessels was evaluated as previously described (9–11). The peripheral region of mouse RPE-choroid-sclera pieces were cut into 1 mm x 1 mm-size pieces. Each piece was implanted into Growth Factor Reduced Matrigel (Corning Inc.) in a 24-well plate, covered with EGM-2 Endothelial Growth Medium (Lonza, Walkersville, MD, USA), and incubated at 37°C with 5% $CO_2$. The medium was changed every 48 h. DMSO vehicle, 30 µM Nec-1 and/or 30 µM Z-VAD was added on day 3. Images of choroidal sprouting were captured on day 7 using a microscope Axio Imager M2 (Zeiss). The distance from the edge of the RPE-choroid-sclera to the tip of the choroidal sprouting was measured using Photoshop CS3 software, and the average in four different directions was defined as the extent of choroidal sprouting for the piece of RPE-choroid-sclera.

**BMDM Culture**
Bone marrow cells were collected from mouse femur and tibia. Red blood cells were lysed by $NH_4Cl$ solution (STEMCELL Technologies, Cambridge, MA, USA) for 10 min on ice. Bone marrow cells were suspended in IMDM medium (Thermo Fisher Scientific) with 10% FBS and 10 ng/ml M-CSF (BioLegend) and seeded onto the plates at a concentration of 1.0 x $10^6$/ml. The growth medium was changed on day 3. On day 7, the differentiated macrophages were treated with 10 ng/ml IL-4 (BioLegend) for M2 polarization. On day 8, the medium was changed to new polarization medium with DMSO vehicle, Nec-1, or Nec-1 with Z-VAD at 30 µM. Supernatant and cell samples were collected after 24 or 48 h.

**Flow Cytometry**
Cultured BMDMs were detached using 5 mM EDTA and gentle scraping and suspended in flow cytometry staining buffer (Thermo Fisher Scientific). Fc receptors were blocked by anti-mouse CD16/CD32 antibody (eBioscience). Dead cells were stained using the LIVE/DEAD Fixable Violet Stain Kit (Thermo Fisher Scientific). Then the cells were treated with fixation and permeabilization buffer (eBioscience), and stained with 1 $\mu$l of the original stock solution of the following antibodies: anti-CD68 antibody (FA-11)-eFluor 660 (Thermo Fisher Scientific), anti-CD206 antibody-FITC (BioRad), and anti-CD163 antibody-FITC (Bioss Antibodies, Woburn, MA, USA). Stained cells were processed using a BD LSR II Analyzer (BD Biosciences, Billerica, MA, USA), and the data were analyzed using FlowJo 10.2 software (FlowJo, LLC, Ashland, OR, USA). For analysis, a pulse geometry gate (FSC-A vs FSC-H) was used to select a single cell population, followed by the exclusion of a dead cell population using intracellular violet staining. The selected viable singlet macrophages were plotted through the gating of eFluor 660 vs FITC.

**Statistical Analysis**
Statistical analyses were performed using Prism software (GraphPad, La Jolla, CA, USA). Student's t-test or one-way ANOVA followed by post hoc Tukey's test was used for numerical data, and the Chi-square test was used for categorical data. $P<0.05$ was considered statistically significant.

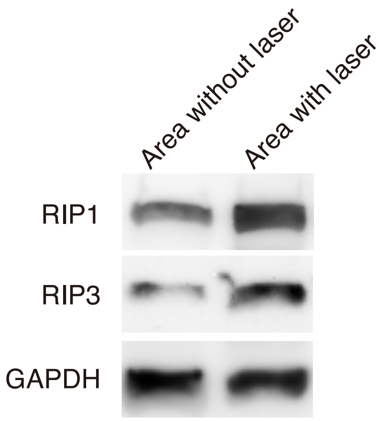


**Fig. S1.** Western blot analysis of RIP1 and RIP3 in RPE-choroid on day 5 after CNV induction. Left lane is the samples dissected from the area without laser-induced CNV lesions, while the right lane is the samples dissected from the area with laser-induced CNV lesions of the same eyes.

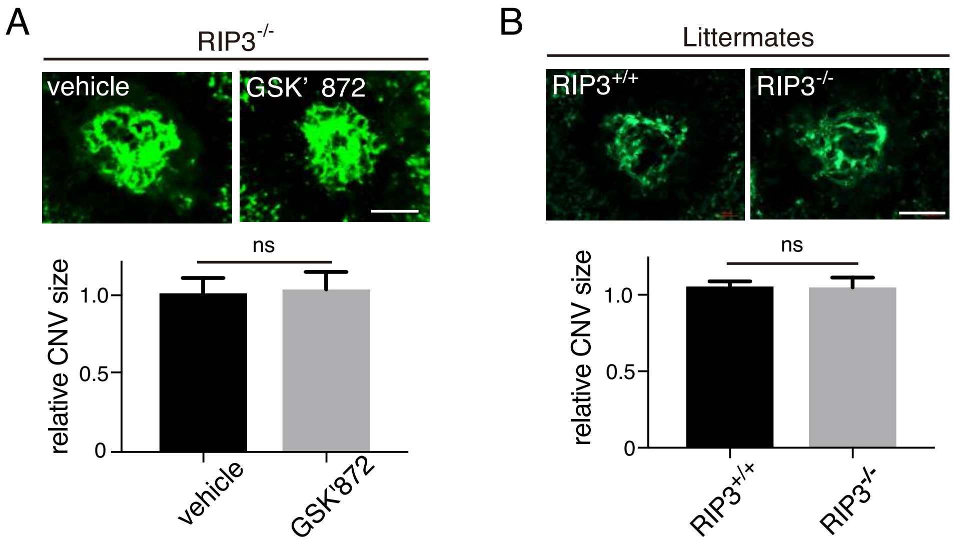


**Fig. S2.** (A) DMSO vehicle or GSK'872 was intravitreally injected to the eyes of $RIP3^{-/-}$ mice after CNV induction (day 0), and CNV size was assesed on day 7. n=6 eyes per group. (B) CNV size on day 7 in $RIP3^{+/+}$ and $RIP3^{-/-}$ mice from the same breeding pairs of $RIP3^{+/-}$ mice. n = 8 eyes per group; scale bar, 100 µm. not significant by Student's t-test. Data are mean ± SEM.

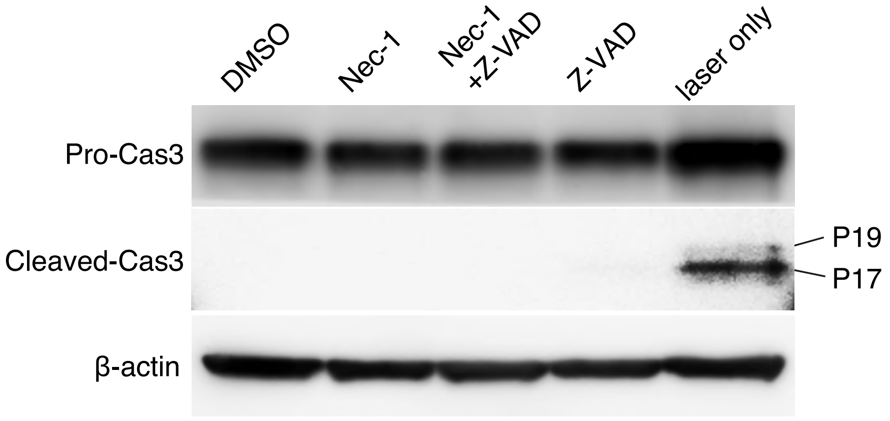


**Fig. S3.** The effects of Nec-1 and/or Z-VAD treatment on caspase-3 activation in non-lasered eyes. Western blot analysis using RPE-choroid samples 4 days after injections with DMSO vehicle, Nec-1, and/or Z-VAD. The laser only lane served as positive control using samples from lasered eyes.

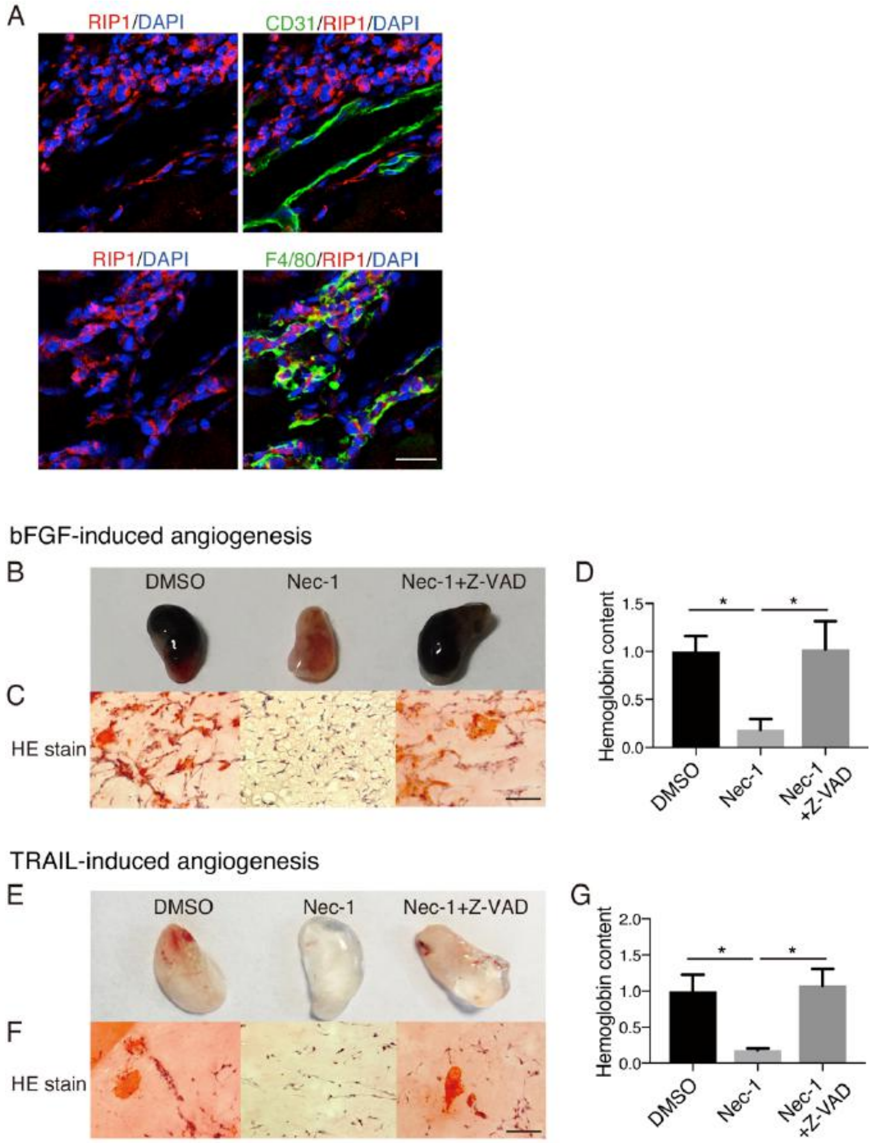


**Fig. S4**. Catalytic inhibition of RIP1 attenuates angiogenesis in Matrigel plugs in vivo through caspase activation. (A) Immunohistochemistry of RIP1 and its localization in F4/80(+) macrophages and CD31(+) endothelial cells in Matrigel plugs on day 7 after implantation. Scale bar, 100 μm. (B–D) in vivo Matrigel plug assay using bFGF (150 ng/ml) with heparin (20 units/ml) activation as the angiogenesis inducer. (E–G) in vivo Matrigel plug assay using TRAIL (400 ng/ml) as the angiogenesis inducer. (B and E) Plugs obtained from WT mice on day 7 after inoculation with Matrigel containing DMSO vehicle, Nec-1 (30 μM), or Nec-1 (30 μM) + Z-VAD (30 μM). (C and F) Representative hematoxylin and eosin (HE) staining images from each treatment group showing neovascularization with RBCs inside. Scale bars, 100 μm (D and G) Relative hemoglobin content normalized to the weight of the corresponding Matrigel plugs containing DMSO vehicle, Nec-1 (30 μM), or Nec-1 (30 μM) + Z-VAD (30 μM). n = 6 plugs per group. *$P < 0.05$; one-way ANOVA and post-hoc Tukey's test (D, G). Data are mean ± SEM.

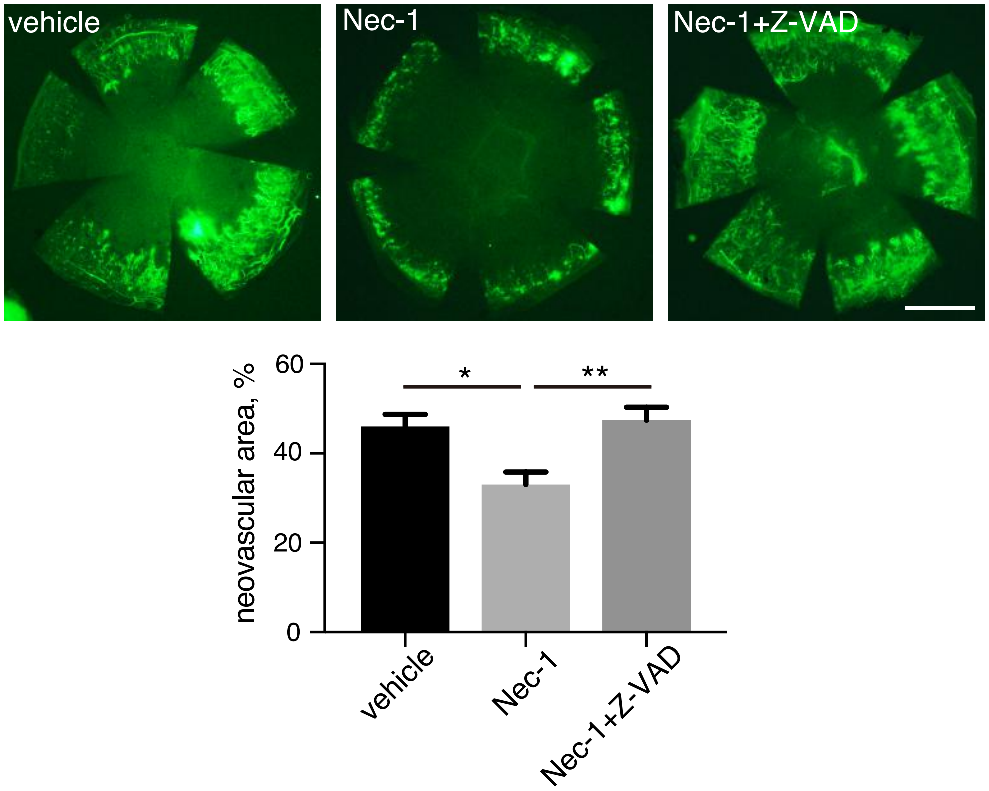


**Fig. S5**. Corneal neovascularization after alkali injury. After alkali injury the WT mice were treated with subconjunctival injections of DMSO vehicle, Nec-1 (300 μM), or Nec-1 (300 μM) + Z-VAD (300 μM). 10 days after injury neovessels were statined with fluorescein concanavalin-A perfusion, and the cornea was flatmounted. Scale bar, 1 mm. n = 6 cornea per group. *$P < 0.05$, **$P < 0.01$; one-way ANOVA and post-hoc Tukey's test. Data are mean ± SEM.

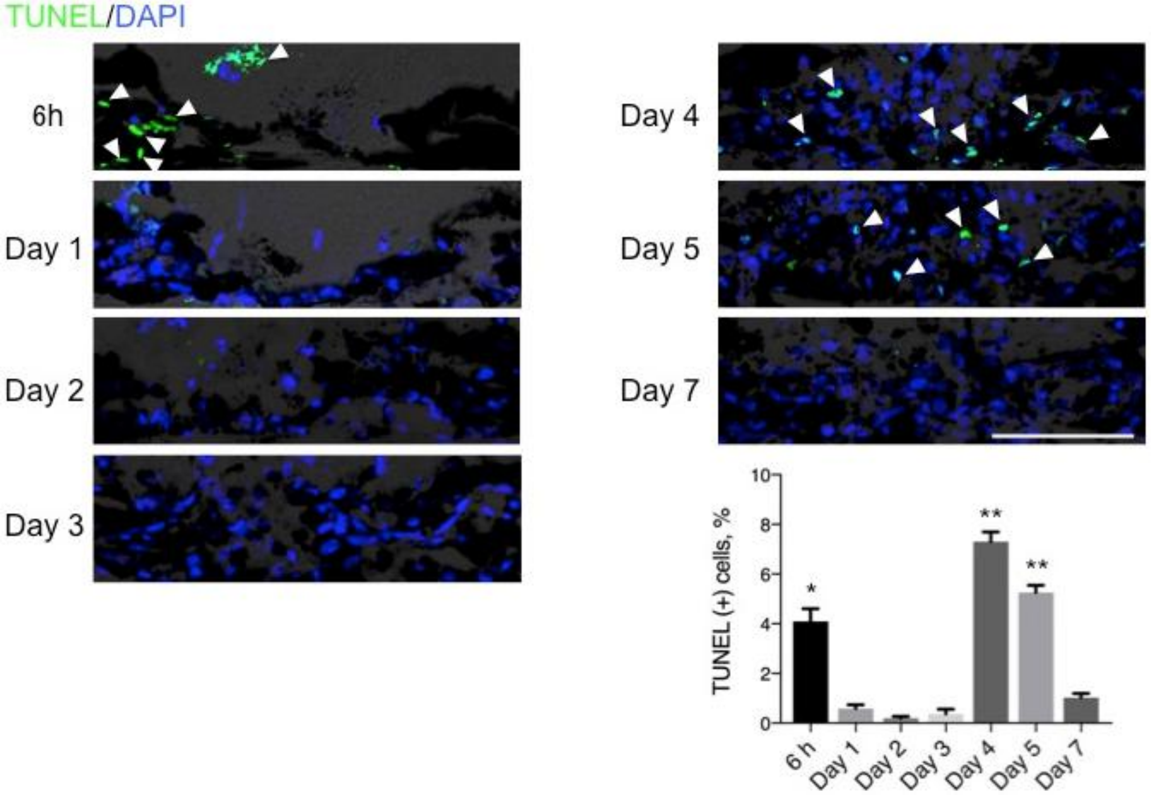


**Fig. S6**. TUNEL staining of sections of CNV lesions during 6h–day7 after CNV induction. **P < 0.01, *P < 0.05; one-way ANOVA and post-hoc Tukey's test. Data are mean ± SEM. Scale bar, 50μm.

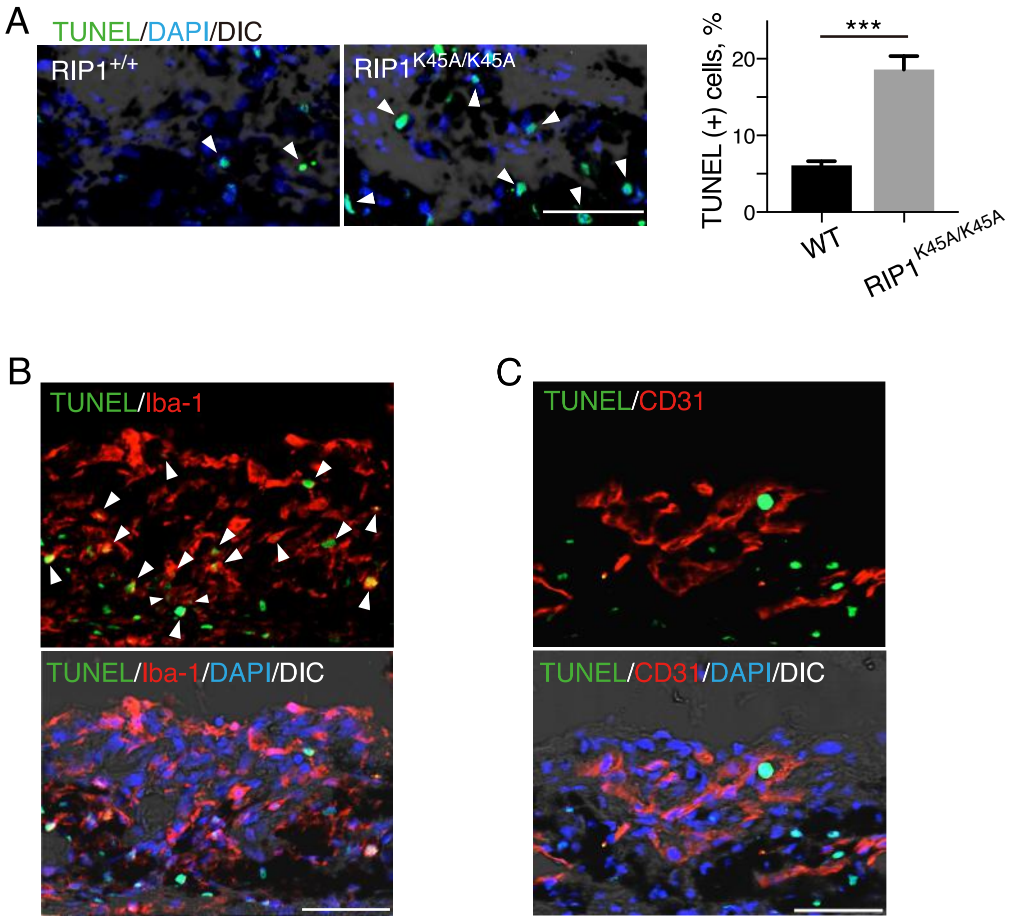


**Fig. S7.** TUNEL staining in Iba1 (+) macrophages (A) TUNEL staining of CNV sections at day 5 after CNV induction in eyes of WT and RIP1$^{K45A/K45A}$ mice. n = 6 lesions per group; scale bar, 50 μm. (B) TUNEL staining and subsequent immunohistochemistry of Iba-1 on CNV sections at day 5 after CNV induction and intravitreal injections with Nec-1. Arrows indicate TUNEL(+) in Iba-1 (+) macrophages/microglia. Scale bar, 50 μm. (C) TUNEL staining and subsequent immunohistochemistry of CD31 on CNV sections at day 5 after CNV induction and intravitreal injections with Nec-1. Scale bar, 50μm.

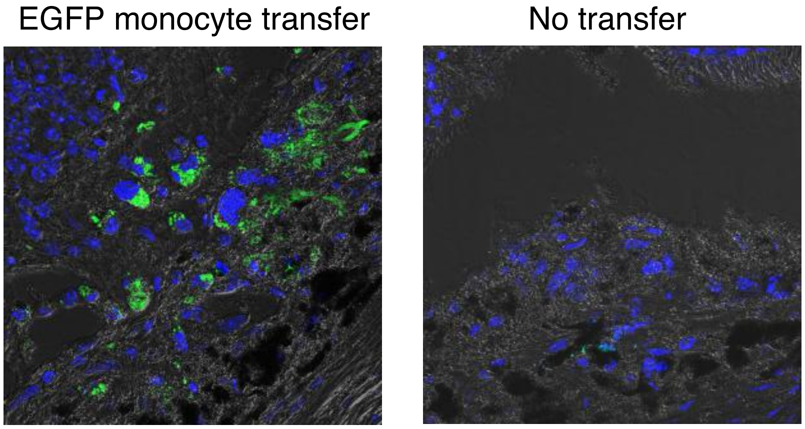


**Fig. S8.** Bone marrow macrophages from EGFP mice transferred to intraperitoneal cavity of WT mice on day 2 after CNV induction. CNV sections were observed on day 4 to validate that transferred macrophages migrate to CNV lesions.

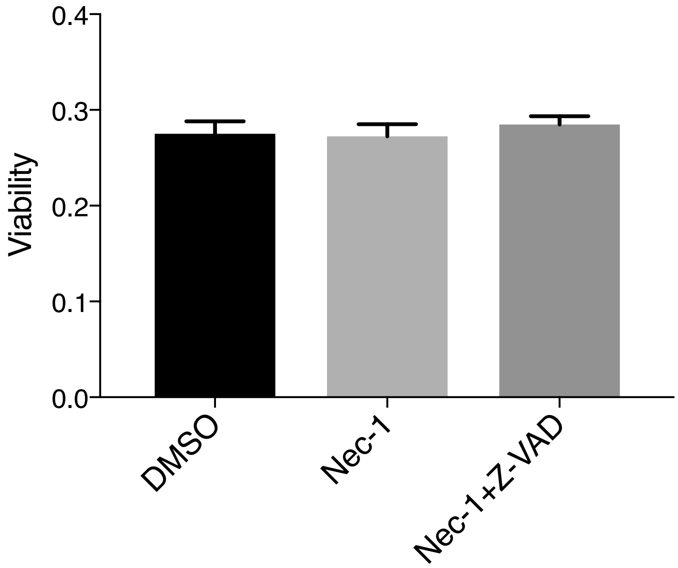


**Fig. S9**. Cellular viability of M2 (IL-4) macrophages is not affected by Nec-1 (30μM) treatment with or without Z-VAD (30 μM) for 48 h. Viability was assessed as dehydrogenase activity of live cells using CCK-8 assay kit. n = 4 per group. Not significant by one-way ANOVA. Data are mean ± SEM.

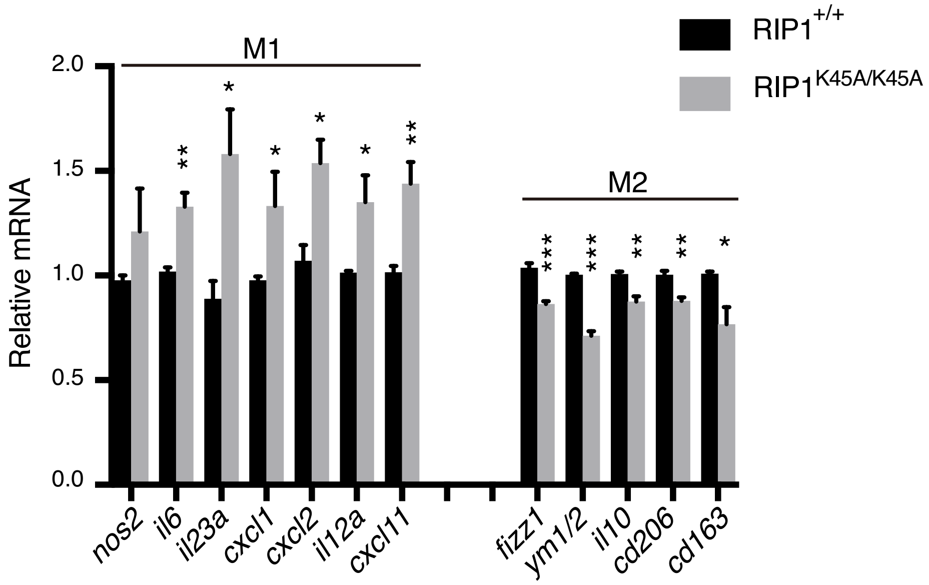


**Fig. S10**. Relative mRNA levels of M2-polarized BMDMs from WT and $RIP1^{K45A/K45A}$ mice. n = 3–4 samples per group. *P < 0.05, **P < 0.01, ***P < 0.001; Student's t-test. Data are mean ± SEM.

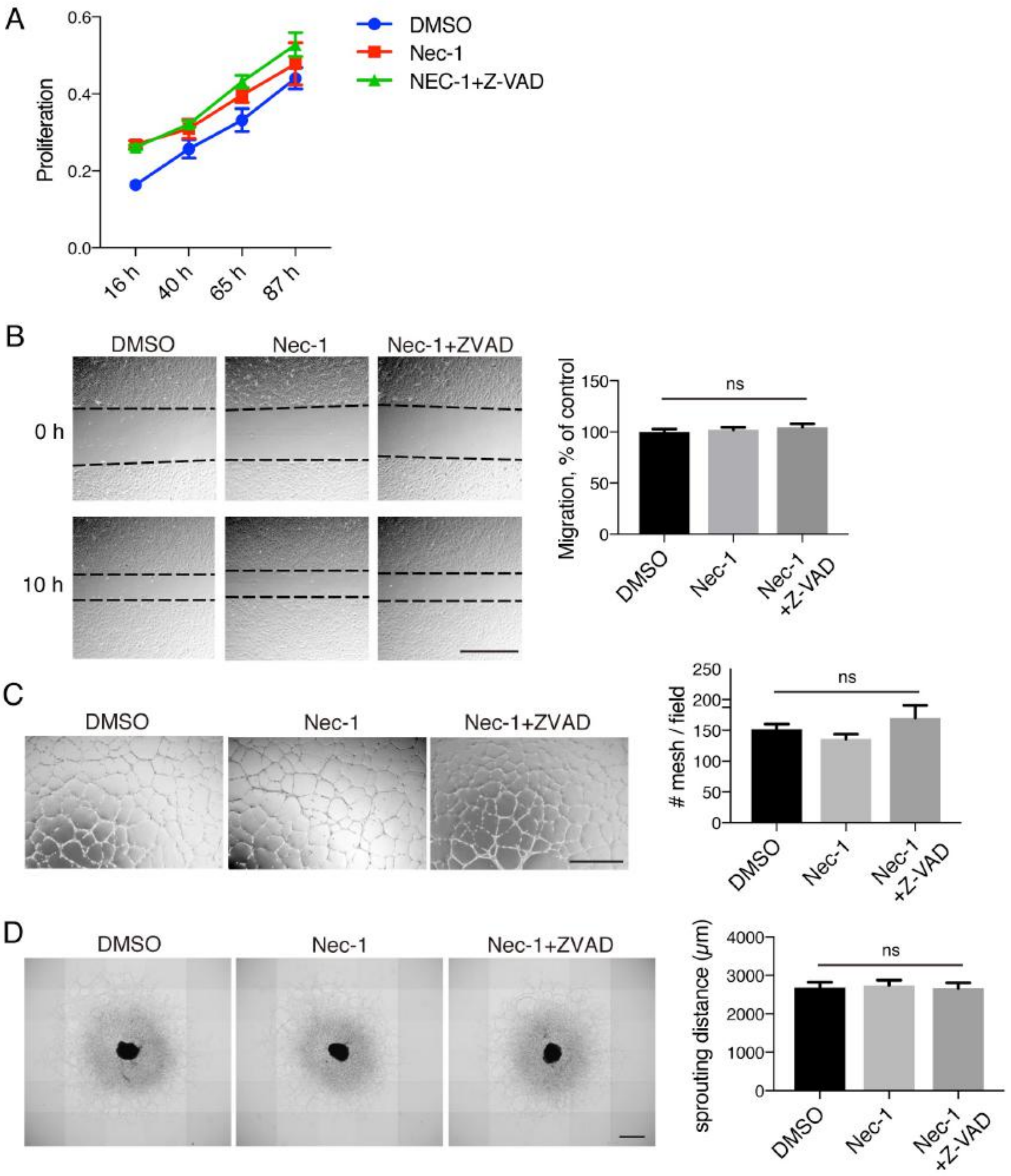


**Fig. S11**. Endothelial cells are not affected by catalytic inhibition of RIP1. (A) Proliferation of HUVEC treated with DMSO vehicle, Nec-1 (30 μM), or Nec-1 (30 μM) + Z-VAD (30 μM). n = 4 per group. *P < 0.05 for Nec-1+Z-VAD at 87 h compared to DMSO control. (B) Migration during 10 h was assessed in HUVEC treated with DMSO vehicle, Nec-1 (30 μM), or Nec-1 (30 μM) + Z-VAD (30 μM). n = 4 per group; scale bar, 1 mm. (C) In vitro tube formation assay of HUVEC treated with DMSO vehicle, Nec-1 (30 μM), or Nec-1 (30 μM) + Z-VAD (30 μM). n = 4 per group; scale bar, 1 mm. (D) Ex vivo choroidal sprouting after 4 days of treatment with DMSO vehicle, Nec-1 (30 μM), or Nec-1 (30 μM) + Z-VAD (30 μM). n = 4 per group; scale bar, 1 mm. ns, not significant; one-way ANOVA (A–D). Data are mean ± SEM.

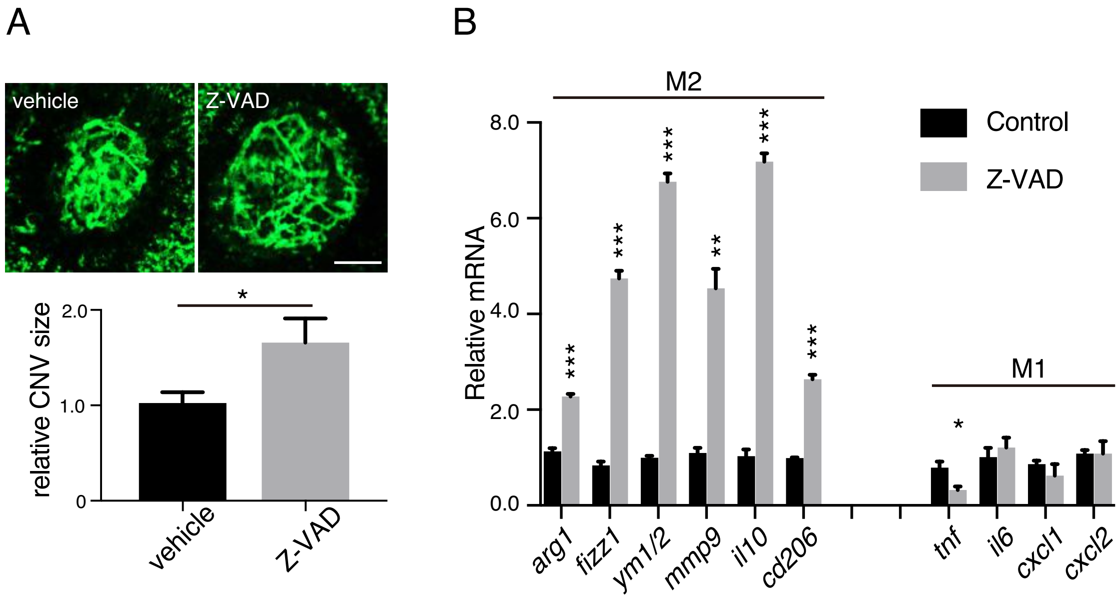


**Fig. S12**. (A) DMSO vehicle or Z-VAD was intravitreally injected to the eyes of WT mice after CNV induction (day 0), and CNV size was assesed on day 7. n=10 eyes per group; scale bar, 100 µm. *P < 0.05 by Student's t-test. Data are mean ± SEM. (B) Relative mRNA levels of M2-polarized BMDM treated with DMSO vehicle or Z-VAD (10 µM) for 24 h. n = 3 samples per group. *P < 0.05, **P < 0.01, ***P < 0.001; Student's t-test. Data are mean ± SEM

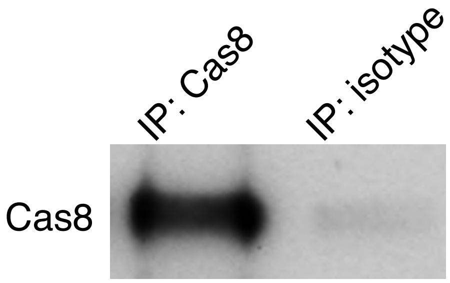


**Fig. S13.** Validation test for the specificity of the caspase-8 antibody used for immunoprecipitation (IP) in the present study. RPE-choloid lysate from lasered mouse eyes were immunoprecipitated using mouse anti-mouse caspase-8 or isotype control antibody, then immunobloted using a rabbit anti-mouse caspase-8 antibody.

**Table S1**. Antibodies for immunofluorescence and western blot used in this study.

| Antigen | Company | Catalogue # | Dilution (immunofluorescence) | Dilution (western or IP) |
|---|---|---|---|---|
| RIP1 | BD Biosciences | 610458 | | 1:1000 |
| RIP1 | CST | 3493 | 1:200 | |
| RIP1 (for human) | Abcam | ab72139 | 1:300 | |
| CD68 (for human) | Abcam | ab955 | 1:200 | |
| RIP3 | NOVUS | NBP1-77299 | | 1;1000 |
| CD31 | BD Pharmingen | 550274 | 1:20 | |
| Iba1 | Wako | 019-19741 | 1:1000 | |
| F4/80 | BIO-RAD | MCA497 | 1:100 | |
| Cleaved Caspase-3 | CST | 9664 | 1:1200 | 1:1000 |
| Caspase-3 | CST | 9662 | | 1:1000 |
| Cleaved Caspase-8 | CST | 8592 | | 1:1000 |
| Caspase-8 | CST | 4927 | | 1:1000 |
| Caspase-8 | Enzo | ALX-804-447 -C100 | | 1:200 (IP) |
| FADD | Santa Cruz | sc-271748 | | 1:200 |
| FADD | Millipore | 05-486 | | 1:1000 |
| pSTAT6 (Tyr641) | Thermo Fisher | 700247 | | 1:1000 |
| STAT6 | CST | 9362 | | 1:1000 |
| pSTAT1 (Tyr701) | CST | 9167 | | 1:1000 |
| STAT1 | CST | 9172 | | 1:1000 |
| CD206 | Abcam | ab64693 | 1:1000 | |
| GAPDH | CST | 2118 | | 1:1000 |
| β-actin | abcam | ab8227 | | 1:3000 |

**Table S2**. Primer sequences.

| Genes (Mouse) | Sequences (5'-3') |
|---|---|
| GAPDH | Forward: CACATTGGGGGTAGGAACAC<br>Reverse: AACTTTGGCATTGTGGAAGG |
| ARG1 | Forward: GTGAAGAACCCACGGTCTGT<br>Reverse: CTGGTTGTCAGGGGAGTGTT |
| Fizz1 | Forward: TGCTGGGATGACTGCTACTG<br>Reverse: CTGGGTTCTCCACCTCTTCA |
| Ym1/2 | Forward: GGGCATACCTTTATCCTGAG<br>Reverse: CCACTGAAGTCATCCATGTC |
| MMP9 | Forward: CGTCGTGATCCCCACTTACT<br>Reverse: AACACACAGGGTTTGCCTTC |
| MMP12 | Forward: TTTCTTCCATATGGCCAAGC<br>Reverse: GGTCAAAGACAGCTGCATCA |
| IL10 | Forward: CCAAGCCTTATCGGAAATGA<br>Reverse: TTTTCACAGGGGAGAAATCG |
| PDGFB | Forward: AGCAGAGCCTGCTGTAATCG<br>Reverse: GGCTTCTTTCGCACAATCTC |
| WNT5A | Forward: CAAATAGGCAGCCGAGAGAC<br>Reverse: CTCTAGCGTCCACGAACTCC |
| WNT7B | Forward: TACTACAACCAGGCGGAAGG<br>Reverse: GTGGTCCAGCAAGTTTTGGT |
| IL4 | Forward: TCAACCCCCAGCTAGTTGTC<br>Reverse: TGTTCTTCGTTGCTGTGAGG |
| IL13 | Forward: CAGCTCCCTGGTTCTCTCAC<br>Reverse: CCACACTCCATACCATGCTG |
| iNOS | Forward: CACCTTGGAGTTCACCCAGT<br>Reverse: ACCACTCGTACTTGGGATGC |
| TNF | Forward: AGTCCGGGCAGGTCTACTTT<br>Reverse: GGTCACTGTCCCAGCATCTT |
| IL6 | Forward: AGTTGCCTTCTTGGGACTGA<br>Reverse: CAGAATTGCCATTGCACAAC |
| IL18 | Forward: GCCTCAAACCTTCCAAATCA<br>Reverse: GTGAAGTCGGCCAAAGTTGT |
| IL23A | Forward: GACTCAGCCAACTCCTCCAG<br>Reverse: GGCACTAAGGGCTCAGTCAG |
| CXCL1 | Forward: GCTGGGATTCACCTCAAGAA<br>Reverse: TCTCCGTTACTTGGGGACAC |
| CXCL2 | Forward: AAGTTTGCCTTGACCCTGAA<br>Reverse: AGGCACATCAGGTACGATCC |
| CXCL10 | Forward: CCCACGTGTTGAGATCATTG<br>Reverse: CACTGGGTAAAGGGGAGTGA |
| CXCL11 | Forward: CGGCTGCGACAAAGTTGAAG<br>Reverse: GCATGTTCCAAGACAGCAGA |
